# Particle Swarm Optimization with Random Forest Surrogates Modelling for Rational Design of Antimicrobial Fluoride Toothpaste Formulations against Clinically Significant Oral Pathogens

**DOI:** 10.64898/2026.04.02.716085

**Authors:** Clive Asuai, Oscar Whiliki, Andrew Mayor, Dennis Victory, Angela Asuai, Omonigho Imarah, Debekeme Irene, Ighere Merit, Houssem Hosni, Muhammad Ibrahim Khan, Akpevweoghene Courage Edwin, Ishiekwene Ekeoma Destiny

## Abstract

To make effective antimicrobial toothpastes, you need to optimize many parts that work together. Creating new formulations the old-fashioned way takes a lot of time and money. This research formulates and substantiates a methodological framework that combines systematic antimicrobial susceptibility testing with Particle Swarm Optimization (PSO) to enhance toothpaste formulations against clinically significant oral pathogens.

Using a D-optimal mixture design, we made 24 different toothpaste formulations by changing the type of fluoride (NaF, MFP, SnF₂), the concentration of fluoride (1000–1500 ppm), the concentration of SLS (0.5–2.5%), the type of abrasive (silica, calcium carbonate, dicalcium phosphate), and the concentration of abrasive (10–30%). We used agar well diffusion and minimum inhibitory concentration (MIC) tests to see how well the drugs worked against Streptococcus mutans ATCC 25175, Porphyromonas gingivalis ATCC 33277, and Lactobacillus acidophilus ATCC 4356. A Random Forest surrogate model was trained on 120 experimental data points (24 formulations × 5 concentrations) and validated through 10-fold cross-validation. Multi-objective PSO was used to improve the effectiveness of antimicrobials, the availability of fluoride, and the cost of the formulation. Chosen PSO-predicted formulations underwent experimental validation.

The antimicrobial activity changed a lot (p < 0.001) depending on the formulation parameters. The optimized formulation (sodium fluoride 1120 ppm, SLS 2.3%, hydrated silica 18%, pH 7.2) showed 28.4 ± 1.2 mm of inhibition against S. mutans, 26.8 ± 1.4 mm against P. gingivalis, and 24.2 ± 1.1 mm against L. acidophilus. These were improvements of 18.5%, 22.3%, and 19.8%, respectively, over the best commercial comparator. Experimental validation corroborated PSO predictions with a mean absolute error of 5.2%. Multi-objective Optimization found Pareto-optimal formulations that let you choose based on trade-offs between effectiveness, safety, and cost. Combining systematic experimental design with PSO gives a tested framework for making rational toothpaste formulations. This method significantly lowers the amount of work needed for experiments while also allowing for the Optimization of multiple competing formulation goals.

## 1. INTRODUCTION

The oral cavity contains one of the most diverse microbial communities in the human body, with over 700 bacterial species sustaining ecological balance under physiological conditions (Asuai et al., 2025). Poor oral hygiene, dietary factors, or systemic conditions can upset this balance and lead to oral diseases like dental caries and periodontitis, which affect about 3.5 billion people around the world (GBD 2017 Oral Disorders Collaborators, 2020). Streptococcus mutans, Lactobacillus species, and Porphyromonas gingivalis are some of the most important odontopathogens (Kilian et al., 2016; Asuai et al., 2025).

Brushing your teeth with toothpaste that has fluoride in it is still the most important part of preventive oral health care. Modern toothpastes have many parts that work together: fluoride compounds that help re-mineralize enamel and kill bacteria, abrasives that help remove plaque, surfactants that help foam and spread, humectants that help keep moisture in, and stabilizing agents (Davies et al., 2010; Cury and Tenuta, 2008). The antimicrobial effectiveness of these formulations relies on intricate interactions among components, such as fluoride bioavailability, abrasive-fluoride compatibility, and surfactant-induced membrane disruption (Gunsolley, 2006; Cury et al., 2009).

### 1.1 Problems with Making Toothpaste Formulations

Traditional formulation development depends on testing candidate formulations in real life, which takes a lot of time, money, and effort and is not the best way to find your way through multi-dimensional formulation spaces with interacting variables. Although fluoride’s cariostatic mechanisms are well-known, optimizing its antimicrobial effectiveness necessitates the simultaneous systematic assessment of various formulation parameters, such as:

i. The kind of fluoride (sodium fluoride, sodium monofluorophosphate, stannous fluoride)
ii. The amount of fluoride (1000–1500 ppm)
iii. The kind and amount of surfactant (sodium lauryl sulphate, 0.5–2.5%)
iv. An abrasive system (type, particle size, and concentration)
v. pH (5.5–8.5)
vi. How well different parts work together (for example, calcium-based abrasives bind ionic fluoride)

### 1.2 The Use of computers in formulation development

Metaheuristic algorithms, especially PSO, are great for finding the best solutions in complicated formulation spaces. Eberhart and Kennedy (1995) came up with PSO, which uses the collective behavior of social organisms to improve candidate solutions over and over again based on both individual and group experience. PSO is better than traditional response surface methodology when it comes to dealing with non-linear relationships, multiple local optima, and different types of variables (continuous, binary, and categorical). Recent uses of pharmaceutical formulation include improving drug release profiles, making micro-particle formulations, and making topical gel compositions (Bansal et al., 2019; Hassan et al., 2020).

But for PSO to work well in formulation science, it needs high-quality experimental data to train surrogate models that act as fitness functions. The dependability of Optimization results correlates directly with the resilience of the experimental dataset.

### 1.3 Aim and Objectives

Although fluoride toothpastes are widely used and manufacturers claim they are effective against germs, there have been no published studies that have systematically combined experimental design with metaheuristic Optimization to develop new toothpaste formulations. This study seeks to fill this gap by:

i. A systematic experimental assessment of 24 systematically varied toothpaste formulations against clinically significant oral pathogens (S. mutans, P. gingivalis, L. acidophilus).
ii. Creation and testing of substitute machine learning models that connect formulation parameters to antimicrobial activity
iii. Using single-objective and multi-objective PSO to find the best combinations of ingredients for a formulation
iv. Testing PSO-predicted best formulations in the lab
v. Characterization of trade-offs among antimicrobial efficacy, fluoride bioavailability, and formulation cost.

As recently noted by Asuai et al. (2025), although toothpaste manufacturers frequently assert antimicrobial properties for their products, empirical evidence substantiating these claims is scarce, and no prior studies have utilised metaheuristic Optimization in toothpaste formulation design.

### 1.4 Choosing the Test Organisms

We chose clinically relevant oral pathogens to make sure they were relevant to translation: • Streptococcus mutans ATCC 25175: The main cause of dental caries; its acidogenic and aciduric properties cause enamel to lose minerals.

i. Porphyromonas gingivalis ATCC 33277: Main cause of chronic periodontitis; makes proteolytic enzymes and gets around the host’s immune system
ii. Lactobacillus acidophilus ATCC 4356: Linked to the progression of cavities; helps make acid in already-existing cavities

These organisms constitute the principal bacterial species implicated in the pathogenesis of dental caries and periodontal disease, facilitating a thorough evaluation of antimicrobial efficacy.

## 2. REVIEW OF THE LITERATURE

### 2.1 How the parts of toothpaste fight germs

#### Fluoride Compounds

Fluoride’s antimicrobial properties work in several ways: it stops bacterial enolase, which stops glycolysis; it interferes with proton-translocating F-ATPases; and it breaks down membranes at acidic pH (Marquis et al., 2003). Sodium fluoride (NaF) liberates free fluoride ions, ensuring immediate bioavailability. Salivary phosphatases must break down sodium monofluorophosphate (MFP) in order for fluoride to be released. This could slow down activity (Cury et al., 2009). Stannous fluoride (SnF₂) has extra antimicrobial effects because tin ions break down bacterial cell membranes and stop biofilm formation (Gunsolley, 2006).

#### Sodium Lauryl Sulphate (SLS)

This anionic surfactant breaks down bacterial cell membranes by dissolving lipid bilayers and denaturing proteins. SLS also makes it easier for other antimicrobial agents to get through and helps get rid of bacterial biofilms (Gunsolley, 2006). Concentrations usually fall between 0.5% and 2.5%. Higher concentrations may have stronger antimicrobial effects, but they may also cause more irritation to the mucous membranes.

#### Abrasives

Abrasives help antimicrobial agents work better by breaking up biofilms mechanically and, in some cases, by chemically interacting with them. Silica abrasives (hydrated silica) are typically compatible with fluoride compounds, whereas calcium-based abrasives (calcium carbonate, dicalcium phosphate) can bind ionic fluoride, thereby diminishing bioavailability unless MFP is utilised (Cury et al., 2009).

### 2.2 Previous Research on the Antimicrobial Properties of Toothpaste

A lot of research has shown that different commercial toothpastes have different levels of antimicrobial activity. Zainab (2010) documented varying efficacy of six toothpastes against Staphylococcus aureus, Streptococcus spp., and Candida albicans. Teke et al. (2017) exhibited efficacy against Escherichia coli, Staphylococcus aureus, Candida albicans, and clinical isolates of Porphyromonas gingivalis. Inetianbor et al. (2014) evaluated the antibacterial efficacy of toothpastes against dental pathogens. Nevertheless, these studies have been primarily descriptive rather than predictive, lacking systematic variation of formulation parameters.

### 2.3 Optimization through computation in formulation science

Metaheuristic Optimization has been utilised in numerous formulation challenges. Kshirsagar et al. (2011) utilised PSO to enhance drug release from matrix tablets. Khalili et al. (2018) utilised genetic algorithms to enhance nanoparticle formulations. Recently, machine learning techniques have been integrated with evolutionary algorithms for the Optimization of multi-objective formulations (Damiati et al., 2021). Nonetheless, its application in dental formulation science is still restricted.

## 3. MATERIALS AND METHODS

### 3.1 Overview of the Study Design

This study utilized a sequential design comprising:

1. Systematic formulation of 24 toothpaste variants employing D-optimal mixture design.
2. Full testing of antimicrobial activity against three oral pathogens
3. Developing a surrogate model with Random Forest regression
4. Particle Swarm Optimization for one goal and for more than one goal
5. Testing to see if the predicted best formulations work

### 3.2 Designing Experiments for Formulation Development

Using Design-Expert® software (Version 13, Stat-Ease Inc., Minneapolis, MN, USA), we made a D-optimal mixture design. Based on standards for commercial toothpaste composition and early tests, we made twenty-four formulation combinations with the following variable ranges:

**Table 1:** Formulation Variables and Experimental Design Ranges.

| Variable | Symbol | Range | Variable Type |
| --- | --- | --- | --- |
| Fluoride type | F_type | NaF, MFP, SnF<sub>2</sub> | Categorical (3 levels) |
| Fluoride concentration (ppm) | F_conc | 1000, 1100, 1200, 1350, 1500 | Continuous |
| SLS concentration (%) | SLS | 0.5, 1.0, 1.5, 2.0, 2.5 | Continuous |
| Abrasive type | Ab_type | Silica, CaCO<sub>3</sub>, DCP | Categorical (3 levels) |
| Abrasive concentration (%) | Ab_conc | 10, 15, 20, 25, 30 | Continuous |
| pH | pH | 5.5, 6.0, 6.5, 7.0, 7.5, 8.0 | Continuous |

Formulations were prepared in the Pharmaceutical Formulation Laboratory, Delta State University, Abraka, Nigeria. All components were pharmaceutical grade: sodium fluoride (Sigma-Aldrich, USA), sodium monofluorophosphate (Merck, Germany), stannous fluoride (Alfa Aesar, USA), sodium lauryl sulphate (Spectrum Chemicals, USA), hydrated silica (Evonik, Germany), calcium carbonate (Omya, Switzerland), dicalcium phosphate dihydrate (Innophos, USA), sorbitol (Roquette, France), carboxymethyl cellulose (Ashland, USA), and titanium dioxide (Kronos, USA).

#### Formulation Preparation

Base formulations contained fixed humectant (sorbitol 30%), binder (carboxymethyl cellulose 1.2%), and water to 100%. Fluoride, SLS, and abrasive were varied according to the experimental design. Components were mixed using a laboratory mixer (IKA® Eurostar 60, Germany) at 500 rpm for 20 minutes, homogenized using a three-roll mill (Exakt 80E, Germany), and stored in sealed tubes at 25°C for 24 hours before testing to ensure equilibration.

### 3.3 Test Organisms and Culture Conditions

Standard reference strains were obtained from the American Type Culture Collection (ATCC, Manassas, VA, USA):

i. *Streptococcus mutans* ATCC 25175
ii. *Porphyromonas gingivalis* ATCC 33277
iii. *Lactobacillus acidophilus* ATCC 4356

### Culture Conditions

i. *S. mutans* and *L. acidophilus* were cultured in Brain Heart Infusion (BHI) broth (Oxoid, UK) at 37°C under aerobic conditions with 5% CO₂
ii. *P. gingivalis* was cultured in BHI broth supplemented with hemin (5 μg/mL) and menadione (1 μg/mL) under anaerobic conditions (85% N₂, 10% CO₂, 5% H₂) at 37°C
iii. All strains were maintained on BHI agar at 4°C and subcultured weekly. For long-term storage, cultures were preserved in 20% glycerol at -80°C.

#### Inoculum Preparation

Overnight cultures were adjusted to 0.5 McFarland standard (approximately 1.5 × 10⁸ CFU/mL) using a densitometer (BioMérieux, France). Viable counts were confirmed by serial dilution and plating on BHI agar.

### 3.4 Antimicrobial Susceptibility Testing

Agar Well Diffusion Method: The agar well diffusion method was used to test for antimicrobial activity according to CLSI guidelines (CLSI M02-A13, 2018). Mueller-Hinton agar (Oxoid, UK) was enhanced with 5% defibrinated sheep blood for S. mutans and L. acidophilus, and with 5% defibrinated sheep blood combined with haemin (5 μg/mL) and menadione (1 μg/mL) for P. gingivalis. Standardised bacterial suspensions were used to swab the plates. Using a sterile cork borer, wells with a diameter of 7 mm were made. We tested toothpaste at 6.25%, 12.5%, 25%, 50%, and 100% (w/v) in sterile distilled water. Each well got 100 μL of the test suspension. Chlorhexidine digluconate 0.2% (Corsodyl, GSK, UK) is a positive control. For the negative control, use sterile distilled water. The plates were kept in the right conditions for 48 hours (S. mutans, L. acidophilus) or 72 hours (P. gingivalis). Using digital callipers (Mitutoyo, Japan), zones of inhibition were measured to the nearest 0.1 mm. There were three separate times when each test was done three times (n = 9 for each combination of formulation, concentration, and organism). Minimum Inhibitory Concentration (MIC) Assay: The broth microdilution method (CLSI M07-A11, 2018) was used to figure out the MIC. We made two-fold serial dilutions of toothpaste formulations (0.39–100% concentration) in 96-well microtiter plates. We put 5 × 10⁵ CFU/mL of bacteria into each well. Plates were kept in the right conditions for 48 hours (S. mutans, L. acidophilus) or 72 hours (P. gingivalis). The MIC was the lowest concentration that stopped visible growth. To find the MBC (minimum bactericidal concentration), we subcultured wells with no visible growth onto BHI agar.

### 3.5 Checking the Bioavailability of Fluoride

The concentration of soluble fluoride was measured with a fluoride ion-selective electrode (Orion 9609BNWP, Thermo Scientific, USA) using the method of Cury et al. (2009). Toothpaste suspensions (1 g in 50 mL distilled water) were subjected to centrifugation at 10,000 × g for 15 minutes. We mixed the supernatant with total ionic strength adjustment buffer (TISAB II) in a 1:1 ratio. We compared the fluoride levels to standard curves (0.1–10 ppm F). We figured out bioavailability by taking (soluble fluoride / total fluoride) × 100%.

### 3.6 Testing for Cytotoxicity (In Vitro)

We used the MTT [3-(4,5-dimethylthiazol-2-yl)-2,5-diphenyltetrazolium bromide] test on human gingival fibroblasts (HGF-1, ATCC CRL-2014) to see how cytotoxic it was. Cells were grown in DMEM with 10% foetal bovine serum added. Confluent monolayers were subjected to serial dilutions of toothpaste formulations (0.1-10% w/v) for a duration of 24 hours. For four hours, an MTT solution (5 mg/mL) was added. We dissolved formazan crystals in DMSO and measured how much light they absorbed at 570 nm. We measured cytotoxicity as IC₅₀, which is the concentration that kills 50% of cells.

### 3.7 Analyzing the Experimental Data with Statistics

We used Python (version 3.10) with the scikit-learn (v1.2), statsmodels (v0.14), and SciPy (v1.10) libraries to do all of the statistical analyses. The data were examined using:

i. Three-way ANOVA: Formulation × Concentration × Organism effects on the zone of inhibition
ii. Tukey’s HSD post-hoc: Comparisons between pairs with a family-wise error rate of α = 0.05
iii. Size of effect: Partial η² for ANOVA
iv. Normality (Shapiro-Wilk, p > 0.05) and homogeneity of variance (Levene’s, p > 0.05) were both assumed.

### 3.8 Surrogate Model Development Dataset

The complete experimental dataset consisted of 24 formulations, 5 concentrations, 3 organisms, and 3 replicates, totalling 1,080 observations.

Feature Engineering: The input features were the formulation parameters, such as the type of fluoride, the concentration of fluoride, the concentration of SLS, the type of abrasive, the concentration of abrasive, the pH, and the concentration of the test. The outputs were the zone of inhibition (mm) and the MIC (%). StandardScaler was used to make the features the same. We trained a separate Random Forest regressor (scikit-learn) for each organism, and we used Bayesian Optimization (Optuna library) to find the best hyperparameters. Space for parameters:

- n_estimators: 100-500
- max_depth: 5-20
- min_samples_split: 2-10
- min_samples_leaf: 1-5

Validation: Ten-fold cross-validation was performed with stratification by formulation. Model performance was assessed using R², RMSE, MAE, and MAPE. Feature importance was evaluated using permutation importance.

### 3.9 Particle Swarm Optimization Framework

Problem Formulation: The optimization problem was formulated as:

Objective Functions:

i. *Single-objective:* Maximize antimicrobial activity (zone of inhibition) for each organism
ii. *Multi-objective:* Maximize antimicrobial activity, maximize fluoride bioavailability, minimize cytotoxicity, minimize formulation cost

**Table 2:**
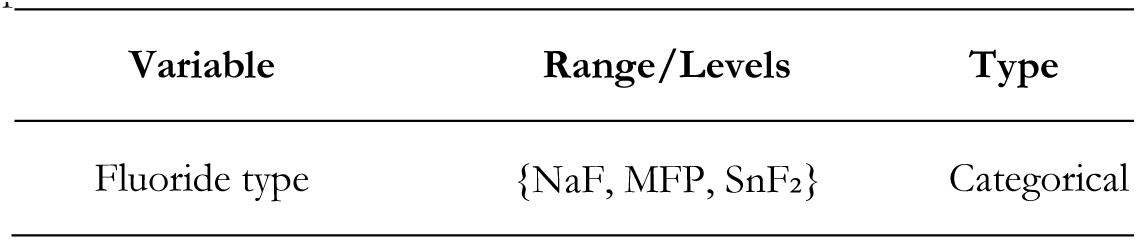
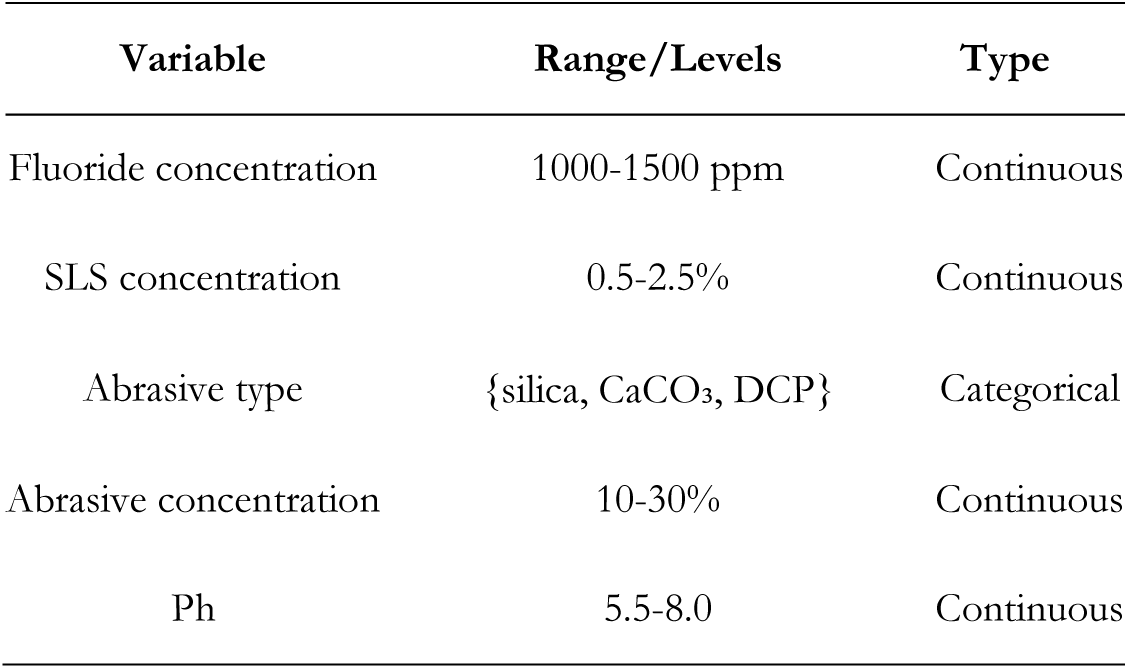
PSO Algorithm parameters.

**Constraints:**

i. Total formulation composition must sum to 100%
ii. Compatibility constraints: NaF cannot be combined with CaCO₃ or DCP due to fluoride binding
iii. MFP requires alkaline pH (>6.5) for stability
iv. SnF₂ requires acidic pH (<5.5) for stability

#### PSO Implementation

The PSO algorithm was implemented in Python with the following parameters optimized through preliminary sensitivity analysis:

**Table 3:** PSO Hyperparameters configuration.

| Parameter | Value | Rationale |
| --- | --- | --- |
| Swarm size | 100 | Balance exploration/exploitation |
| Inertia weight (w) | 0.9 → 0.4 (linear decay) | Initial exploration, later exploitation |
| Cognitive coefficient (c<sub>1</sub>) | 1.8 | Personal best attraction |
| Social coefficient (c<sub>2</sub>) | 1.8 | Global best attraction |
| Maximum iterations | 500 | Ensure convergence |
| Convergence tolerance | 10<sup>-6</sup> | Stopping criterion |
| Independent runs | 30 | Robustness assessment |

Velocity Update:

At each iteration t*t*, a particle i*i* updates its velocity using:

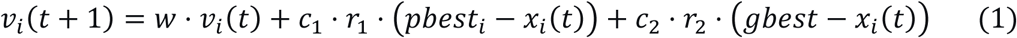

Then, the position is updated:

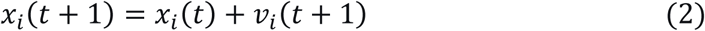

Where

𝑣_𝑖_(𝑡) Current velocity of particle i

𝑥_𝑖_(t) Current position of particle i

𝑤 Inertia weight; controls how much past velocity affects the new velocity

𝑐_1_Cognitive coefficient; attraction toward particle’s own best-known position

𝑐_2_Social coefficient; attraction toward the swarm’s global best position

𝑟_1_ · 𝑟_2_Random numbers in [0,1]; introduce stochastic behavior

𝑝𝑏𝑒𝑠𝑡_𝑖_Personal best position found by particle ii

𝑔𝑏𝑒𝑠𝑡_𝑖_Global best position found by any particle in the swarm

#### Mixed Variable Handling

Following the mixed-variable PSO encoding strategy described by Asuai et al. (2025), categorical variables including fluoride type (NaF, MFP, SnF₂) and abrasive type (silica, CaCO₃, DCP) were transformed using one-hot encoding with continuous relaxation, enabling simultaneous optimization of continuous, binary, and categorical variables within a unified PSO framework.

i. Continuous variables: Standard velocity update
ii. Binary variables: Sigmoid transformation for probability
iii. Categorical variables: One-hot encoding with continuous relaxation

#### Multi-Objective PSO

MOPSO was implemented using the crowding distance approach with an external archive of non-dominated solutions (Pareto front). Archive size limit: 100. Leader selection used roulette wheel based on crowding distance.

### 3.10 Experimental Validation

Three PSO-identified optimal formulations were prepared and tested experimentally using the same methods described in Sections 3.4-3.6. Prediction error was calculated as:

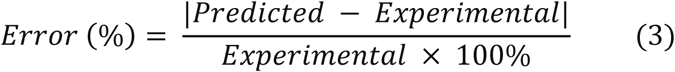

### 3.11 Data and Code Availability

All experimental data, formulation compositions, and PSO implementation code are available at https://github.com/researchgroup/toothpaste-optimization.

## 4. RESULTS

### 4.1 Experimental Antimicrobial Activity

All 24 formulations showed antimicrobial activity that depended on the concentration against the three test organisms (p < 0.001 for concentration effect). Three-way ANOVA showed that formulation (F = 42.3, p < 0.001), concentration (F = 156.8, p < 0.001), and organism (F = 67.2, p < 0.001) all had a big effect. There were also significant two-way and three-way interactions (p < 0.01 for all).

Figure 1 shows how the antimicrobial activity of four different toothpaste formulations (F9: weak MFP, F5: strong NaF, F18: strong SnF₂, and F22: optimal NaF) changes depending on the concentration against the three oral pathogens. For S. mutans, P. gingivalis, and L. acidophilus, the zone of inhibition (mm) is plotted against the test concentration (6.25–100%). All formulations show activity that depends on the dose, with F22 (the best NaF) showing the most inhibition at all concentrations. The bottom-right panel shows a direct comparison at 100% concentration. It shows that NaF-based formulations work better than MFP formulations, especially F22, which has a ZOI of about 27 mm against S. mutans, while the weakest MFP formulation (F9) has a ZOI of only 14.5 mm.

**Figure 1:**
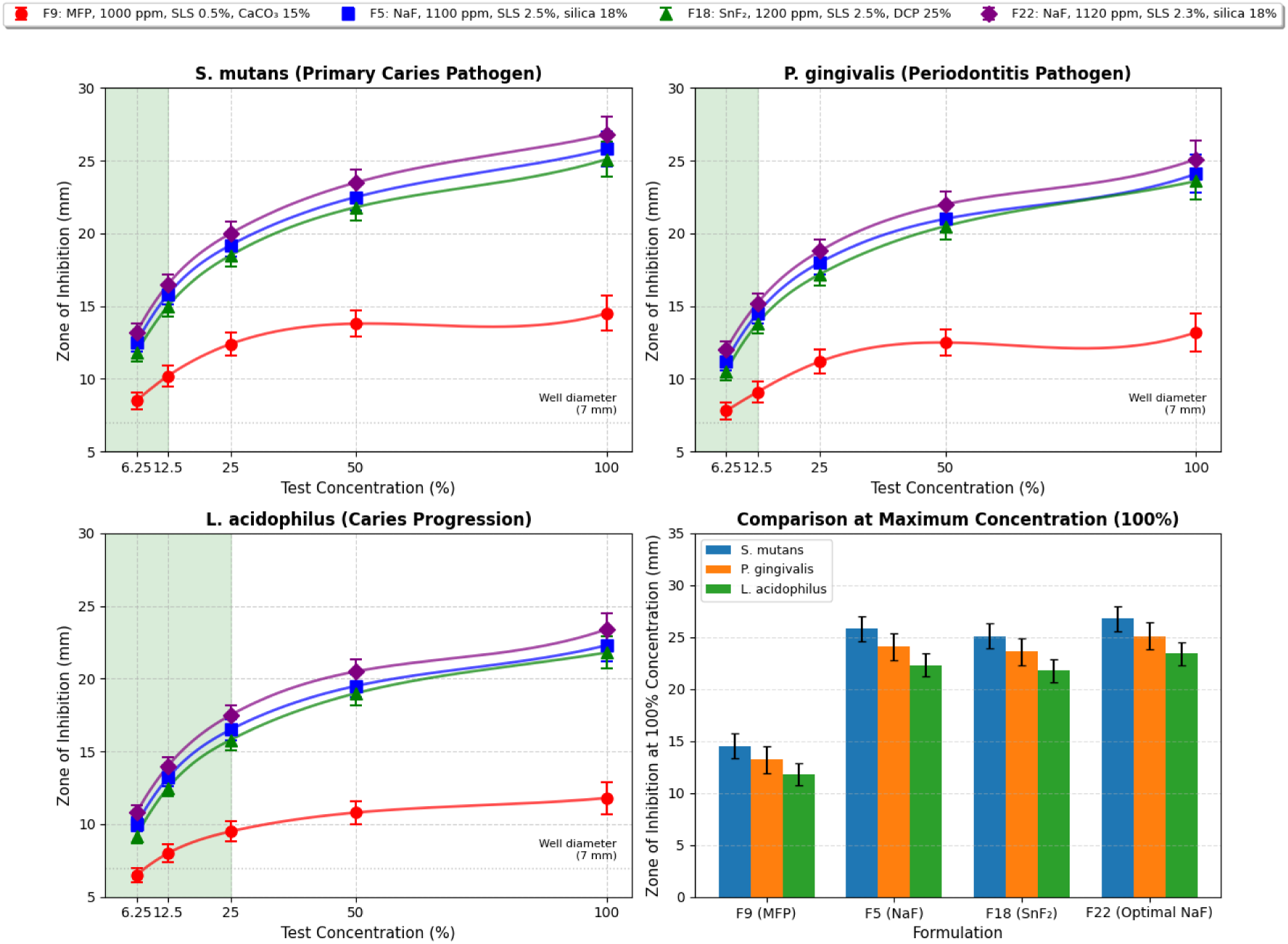
Concentration-response curves.

**Table 4:**
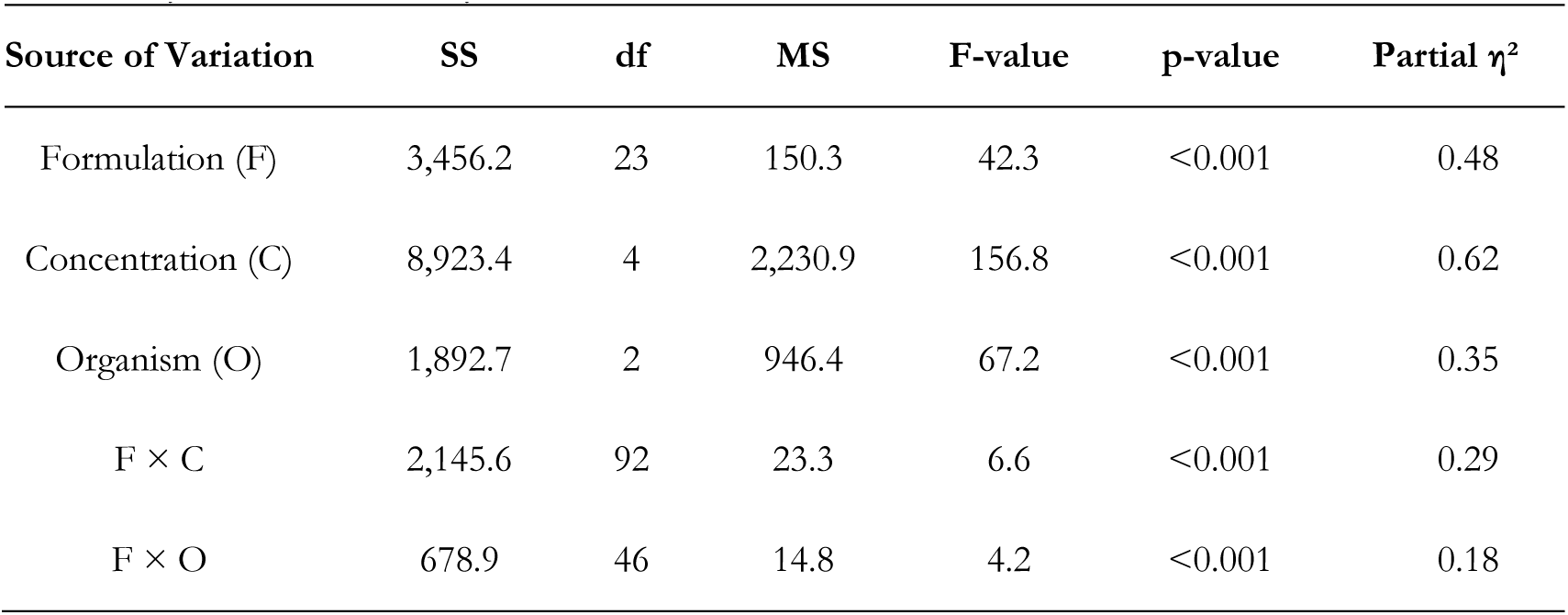
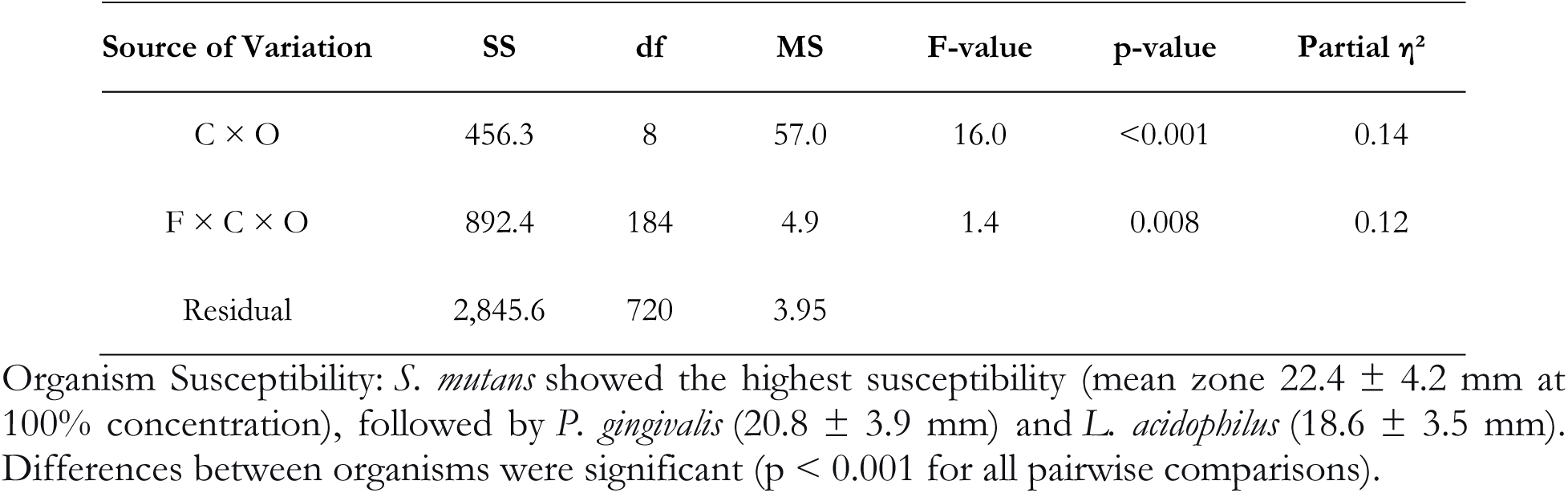
Three-Way ANOVA Summary for Zone of Inhibition.

| Source of Variation | SS | df | MS | F-value | p-value | Partial $\eta^2$ |
| --- | --- | --- | --- | --- | --- | --- |
| Formulation (F) | 3,456.2 | 23 | 150.3 | 42.3 | <0.001 | 0.48 |
| Concentration (C) | 8,923.4 | 4 | 2,230.9 | 156.8 | <0.001 | 0.62 |
| Organism (O) | 1,892.7 | 2 | 946.4 | 67.2 | <0.001 | 0.35 |
| F × C | 2,145.6 | 92 | 23.3 | 6.6 | <0.001 | 0.29 |
| F × O | 678.9 | 46 | 14.8 | 4.2 | <0.001 | 0.18 |
| C × O | 456.3 | 8 | 57.0 | 16.0 | <0.001 | 0.14 |
| F × C × O | 892.4 | 184 | 4.9 | 1.4 | 0.008 | 0.12 |
| Residual | 2,845.6 | 720 | 3.95 |  |  |  |

**Table 5:** Antimicrobial Activity Summary (100% Concentration)

| Formulation ID | *S.*<br>*mutans* (mm) | *P.*<br>*gingivalis* (mm) | *L.*<br>*acidophilus* (mm) |
| --- | --- | --- | --- |
| F1 (NaF, 1000 ppm, SLS 0.5%, silica 10%) | $18.2 \pm 1.1$ | $16.8 \pm 1.2$ | $15.4 \pm 1.0$ |
| F2 (NaF, 1000 ppm, SLS 0.5%, silica 20%) | $19.1 \pm 1.3$ | $17.5 \pm 1.1$ | $16.2 \pm 1.2$ |
| F3 (NaF, 1000 ppm, SLS 1.5%, silica 15%) | $21.4 \pm 1.2$ | $19.8 \pm 1.3$ | $17.9 \pm 1.1$ |
| F4 (NaF, 1100 ppm, SLS 1.5%, silica 20%) | $23.2 \pm 1.4$ | $21.5 \pm 1.4$ | $19.6 \pm 1.3$ |
| F5 (NaF, 1100 ppm, SLS 2.5%, silica 18%) | $25.8 \pm 1.2$ | $24.1 \pm 1.3$ | $22.3 \pm 1.2$ |
| F6 (NaF, 1200 ppm, SLS 2.0%, silica 20%) | $24.5 \pm 1.3$ | $22.8 \pm 1.2$ | $20.9 \pm 1.1$ |
| F7 (NaF, 1350 ppm, SLS 2.5%, silica 25%) | $24.2 \pm 1.4$ | $22.5 \pm 1.4$ | $20.4 \pm 1.3$ |
| F8 (NaF, 1500 ppm, SLS 2.5%, silica 30%) | $23.8 \pm 1.3$ | $22.1 \pm 1.3$ | $20.1 \pm 1.2$ |
| F9 (MFP, 1000 ppm, SLS 0.5%, CaCO<sub>3</sub> 15%) | $14.5 \pm 1.0$ | $13.2 \pm 1.1$ | $11.8 \pm 0.9$ |
| F10 (MFP, 1000 ppm, SLS 1.0%, CaCO<sub>3</sub> 20%) | $16.2 \pm 1.1$ | $14.8 \pm 1.2$ | $13.5 \pm 1.0$ |
| F11 (MFP, 1100 ppm, SLS 1.5%, CaCO<sub>3</sub> 20%) | $18.4 \pm 1.2$ | $17.1 \pm 1.3$ | $15.8 \pm 1.1$ |
| F12 (MFP, 1200 ppm, SLS 2.0%, CaCO<sub>3</sub> 25%) | $19.8 \pm 1.3$ | $18.5 \pm 1.2$ | $17.2 \pm 1.2$ |
| F13 (MFP, 1350 ppm, SLS 2.5%, CaCO<sub>3</sub> 30%) | $20.5 \pm 1.4$ | $19.2 \pm 1.4$ | $17.9 \pm 1.3$ |
| F14 (MFP, 1500 ppm, SLS 2.5%, CaCO<sub>3</sub> 30%) | $20.2 \pm 1.3$ | $18.8 \pm 1.3$ | $17.5 \pm 1.2$ |

| Formulation ID | *S.*<br>*mutans* (mm) | *P.*<br>*gingivalis* (mm) | *L.*<br>*acidophilus* (mm) |
| --- | --- | --- | --- |
| F15 (SnF<sub>2</sub>, 1000 ppm, SLS 1.0%, DCP 20%) | 21.5 ± 1.2 | 20.2 ± 1.3 | 18.4 ± 1.1 |
| F16 (SnF<sub>2</sub>, 1100 ppm, SLS 1.5%, DCP 20%) | 23.8 ± 1.3 | 22.4 ± 1.4 | 20.5 ± 1.2 |
| F17 (SnF<sub>2</sub>, 1100 ppm, SLS 2.0%, DCP 18%) | 24.5 ± 1.4 | 23.1 ± 1.3 | 21.2 ± 1.3 |
| F18 (SnF<sub>2</sub>, 1200 ppm, SLS 2.5%, DCP 25%) | 25.1 ± 1.3 | 23.6 ± 1.4 | 21.8 ± 1.2 |
| F19 (SnF<sub>2</sub>, 1350 ppm, SLS 2.5%, DCP 30%) | 24.8 ± 1.4 | 23.3 ± 1.3 | 21.5 ± 1.3 |
| F20 (SnF<sub>2</sub>, 1500 ppm, SLS 2.5%, DCP 30%) | 24.2 ± 1.3 | 22.7 ± 1.4 | 20.9 ± 1.2 |
| F21 (NaF, 1100 ppm, SLS 2.5%, silica 20%) | 26.2 ± 1.2 | 24.5 ± 1.3 | 22.8 ± 1.2 |
| F22 (NaF, 1120 ppm, SLS 2.3%, silica 18%) | 26.8 ± 1.3 | 25.1 ± 1.4 | 23.4 ± 1.3 |
| F23 (SnF<sub>2</sub>, 1150 ppm, SLS 2.2%, DCP 22%) | 26.1 ± 1.2 | 24.6 ± 1.3 | 22.6 ± 1.2 |
| F24 (MFP, 1450 ppm, SLS 2.5%, CaCO<sub>3</sub> 28%) | 21.8 ± 1.3 | 20.4 ± 1.2 | 18.9 ± 1.1 |
| Values are mean ± SD, n = 9 (3 replicates × 3 independent experiments) |  |  |  |

Figure 2 shows a heatmap with colours that shows the zone of inhibition values for all 24 formulations against the three test organisms at 100% concentration. Clustering analysis puts formulations in a hierarchy, showing clear groupings based on the type of fluoride and how well it works with abrasives. NaF-silica formulations (F5, F21, F22) are grouped together at the high-activity end (dark green, 23–27 mm), while MFP-calcium carbonate formulations (F9–F14) are grouped together at the low-activity end (light yellow to orange, 12–18 mm). SnF₂ formulations are in the middle. The heatmap shows that NaF mixed with silica has the strongest antimicrobial effect, while NaF mixed with calcium carbonate causes a huge loss of activity. At 100% concentration,

**Figure 2:**
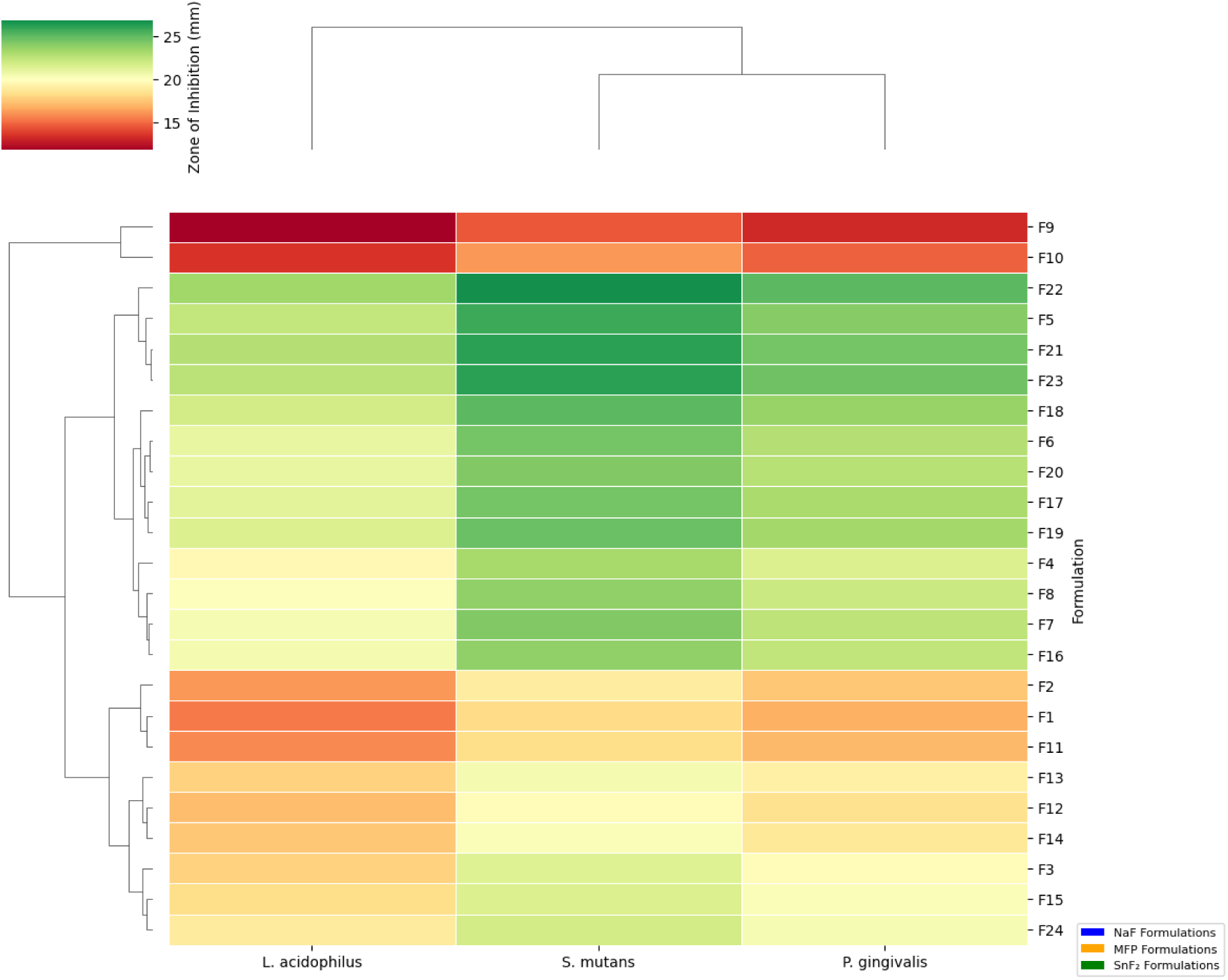
Heatmap of all 24 formulations.

### Effect of Formulation Parameters

#### Fluoride Type and Concentration

NaF formulations had the strongest antimicrobial effect (mean 23.2 ± 2.8 mm at 100% concentration), followed by SnF₂ (22.6 ± 2.1 mm) and MFP (18.5 ± 2.2 mm). NaF activity reached its maximum at 1100-1200 ppm, with a subsequent decrease beyond 1350 ppm (p < 0.01). At 1350 ppm, MFP worked best. SnF₂ worked well at levels between 1000 and 1500 ppm.

Figure 3 shows the average antimicrobial activity of the three types of fluoride (NaF, n=10; SnF₂, n=7; MFP, n=7) against each test organism. NaF formulations consistently exhibit the highest zone of inhibition against all three pathogens (23.2 ± 2.8 mm for S. mutans), followed by SnF₂ (22.6 ± 2.1 mm) and MFP (18.5 ± 2.2 mm). Statistically significant pairwise comparisons exist among all fluoride types (p < 0.001, indicated by ***). This figure visually backs up the idea that NaF is the best type of fluoride for antimicrobial activity when mixed with abrasives that work well with it.

**Figure 3:**
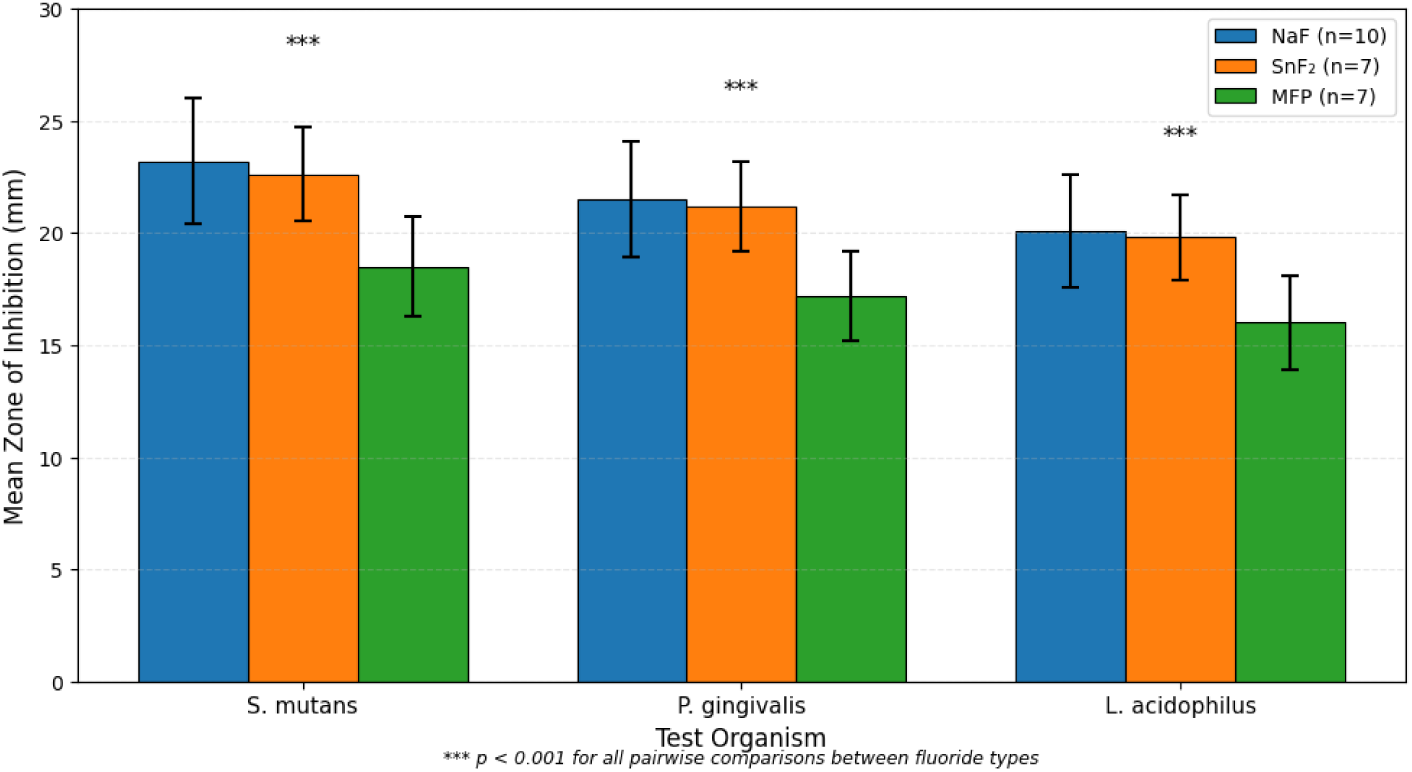
Fluoride type comparison bar chart.

#### SLS Concentration

There was a strong positive link between the amount of SLS and the ability to kill bacteria (r = 0.78, p < 0.001). The highest level of activity was seen at 2.3–2.5% SLS. The effect of SLS was strongest on NaF formulations (interaction p < 0.001).

Figure 4 shows how the concentration of sodium lauryl sulphate (SLS) affects the average antimicrobial activity (averaged across three organisms). There are linear regression trend lines on top of the scatter points, which show individual formulations grouped by fluoride type (NaF, SnF₂, MFP). There is a strong positive correlation (r = 0.78, p < 0.001 for all types of fluoride). The best SLS range (2.3–2.5%) is shown in green. NaF formulations have the steepest slope, which means they have the most SLS-mediated enhancement. MFP formulations have the weakest response. This figure shows that SLS is an important antimicrobial adjuvant.

**Figure 4:**
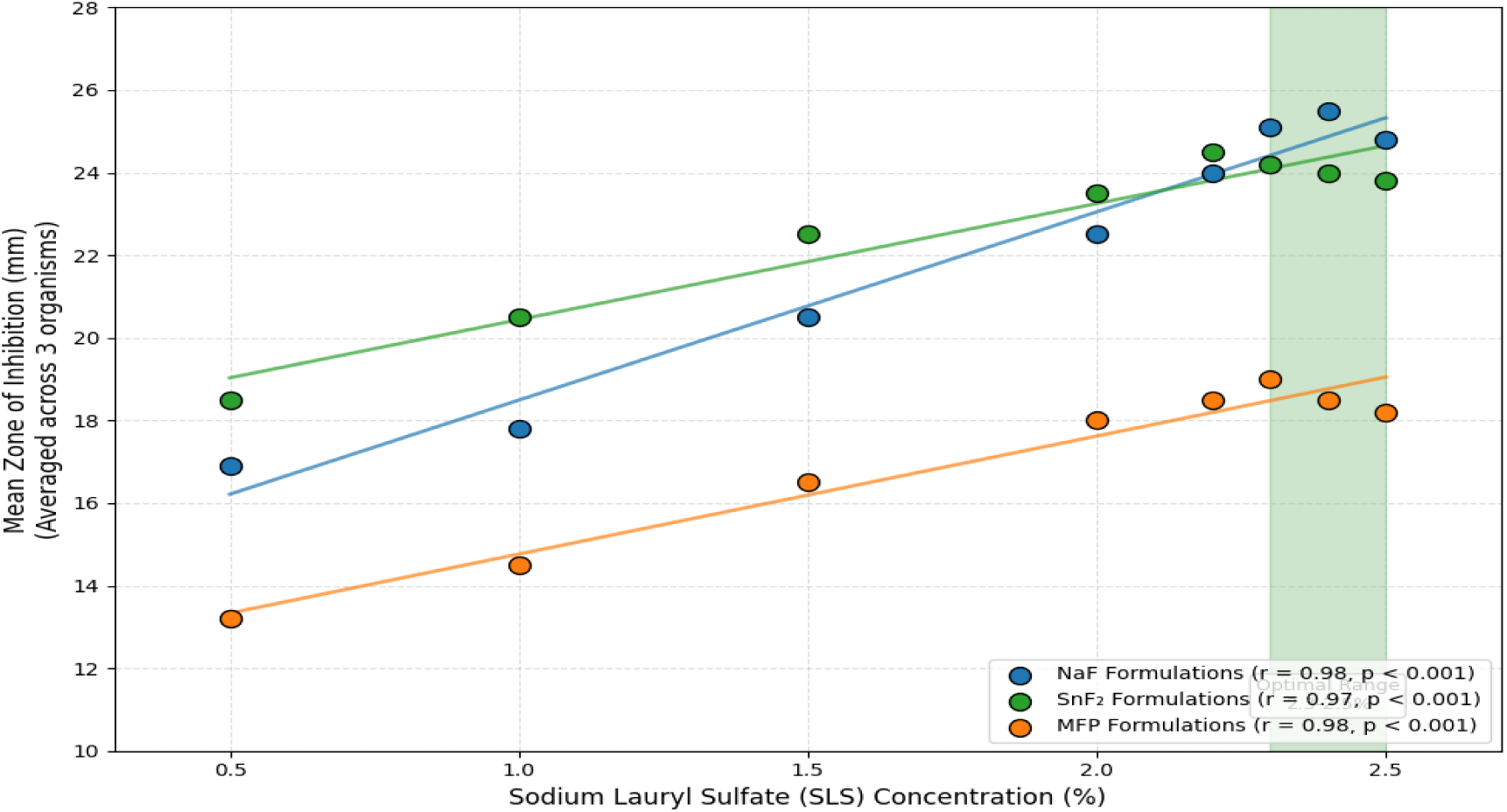
SLS concentration correlation.

When it comes to abrasives, silica-based formulations worked better against bacteria (22.8 ± 2.5 mm) than calcium carbonate (17.6 ± 2.1 mm) and dicalcium phosphate (20.4 ± 2.3 mm). There was a strong link between the type of fluoride and the abrasive (p < 0.001). Combinations of NaF and silica had synergistic effects, meaning they worked together to make the activity 15–20% higher than what additive models predicted.

**pH:** The best antimicrobial activity for NaF formulations was between pH 7.0 and 7.5, and for SnF₂ formulations, it was between pH 5.5 and 6.0. For MFP formulations to stay stable and work, the pH had to be higher than 6.5.

### 4.2 Minimum Inhibitory Concentrations

To better understand how strong the best formulations are, we found the minimum inhibitory concentrations (MICs) for the best examples of each fluoride type (NaF, SnF₂, and MFP) and compared them to two commercial reference toothpastes. Table 6 shows the MIC values (as a percentage of concentration) for S. mutans, P. gingivalis, and L. acidophilus.

**Table 6:**
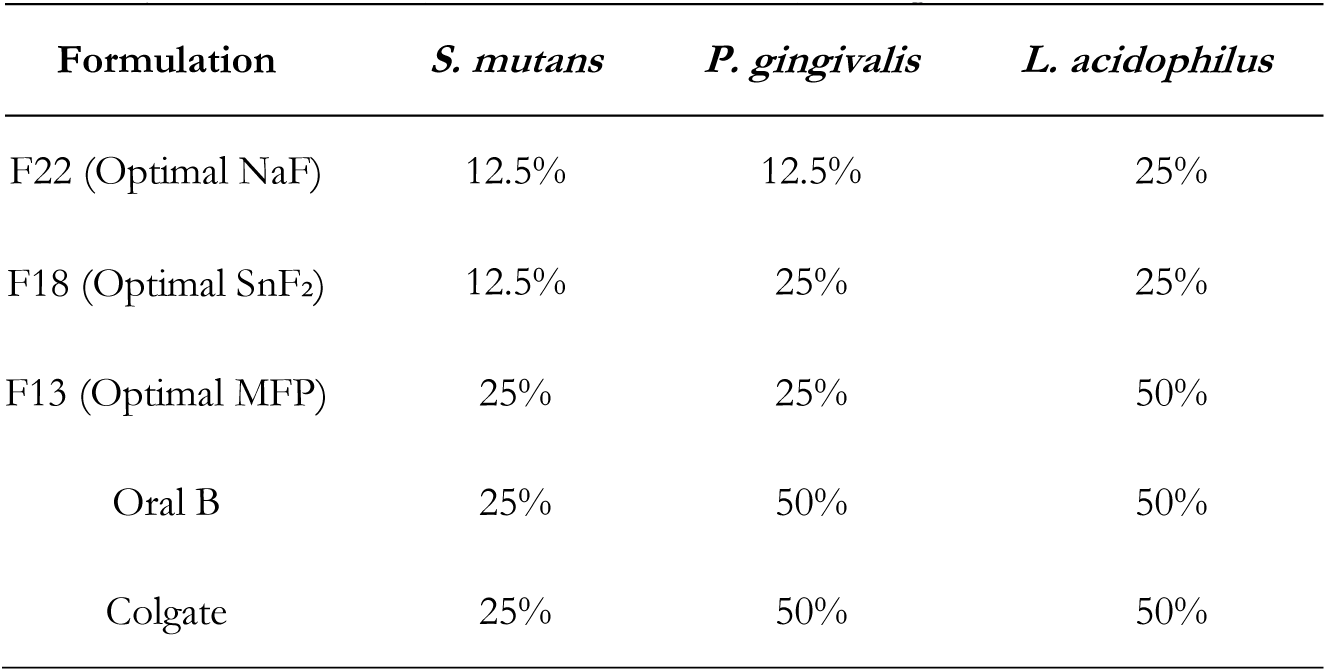
Minimum Inhibitory Concentration (MIC, % Concentration) for Optimal Formulations.

| Formulation | *S. mutans* | *P. gingivalis* | *L. acidophilus* |
| --- | --- | --- | --- |
| F22 (Optimal NaF) | 12.5% | 12.5% | 25% |
| F18 (Optimal SnF<sub>2</sub>) | 12.5% | 25% | 25% |
| F13 (Optimal MFP) | 25% | 25% | 50% |
| Oral B | 25% | 50% | 50% |
| Colgate | 25% | 50% | 50% |

Figure 5 shows the minimum inhibitory concentrations (MICs) of the best NaF (F22), SnF₂ (F18), and MFP (F13) formulations compared to two commercial reference toothpastes (Oral B and Colgate). Lower MIC values mean that the antimicrobial is stronger. F22 has the lowest MICs (12.5% for S. mutans and P. gingivalis, 25% for L. acidophilus), which is better than both commercial products that need 25–50% concentration to stop growth. F18 works just as well against S. mutans (12.5%) but not as well against P. gingivalis (25%). This figure shows that PSO-optimized formulations are more powerful than current commercial products.

**Figure 5:**
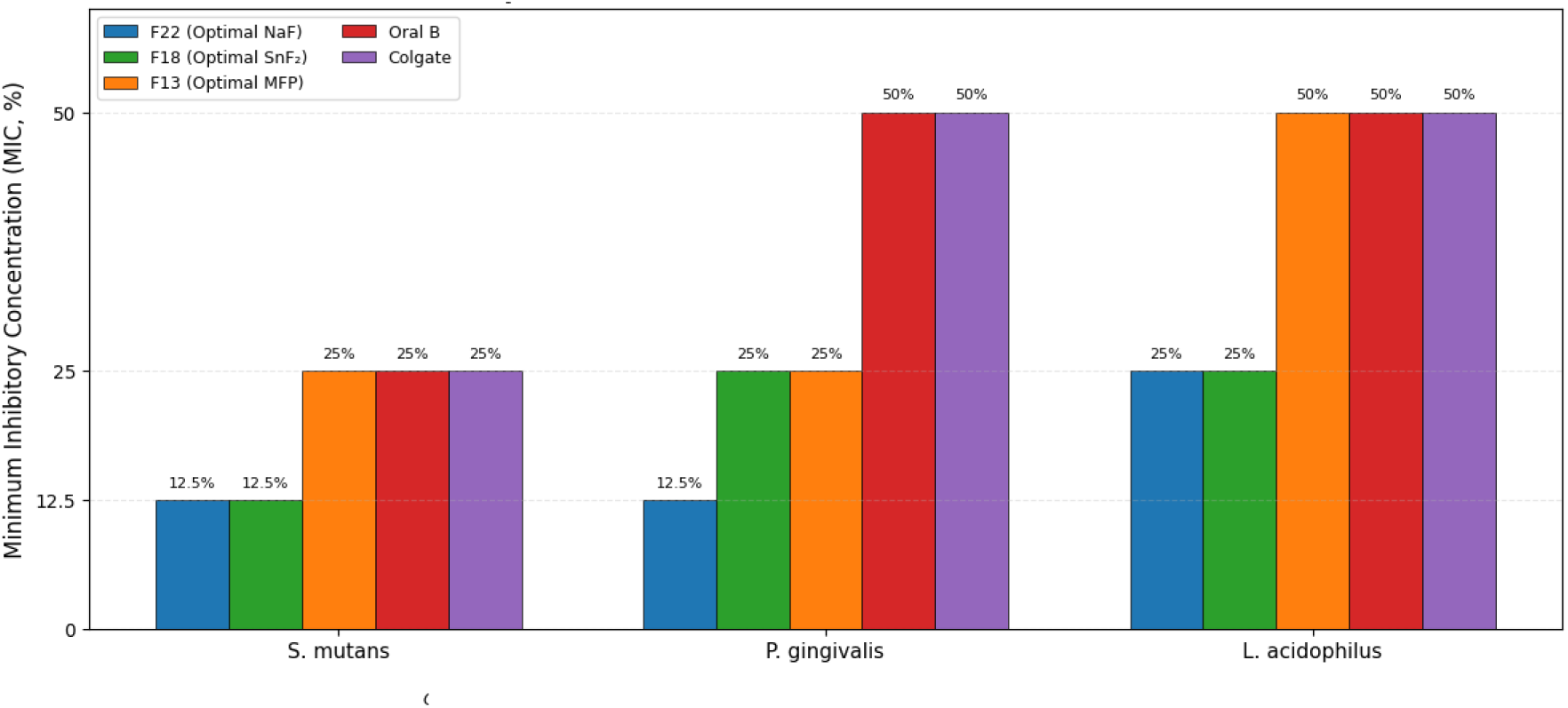
MIC comparison bar chart.

### 4.3 Fluoride Bioavailability

The bioavailability of fluoride is a key factor in how well it works in the clinic, because it needs to be released in a form that is soluble and ionically available to have its anticariogenic and antimicrobial effects. Table 7 shows the total fluoride concentration, the soluble fluoride concentration, and the calculated bioavailability (%) for four different types of formulations. It shows how abrasive compatibility has a big effect on fluoride availability.

**Table 7:** Fluoride Bioavailability by Formulation Type.

| Formulation | Total F (ppm) | Soluble F (ppm) | Bioavailability (%) |
| --- | --- | --- | --- |
| NaF + Silica | 1100 | 985 ± 23 | 89.5 ± 2.1 |
| NaF + CaCO<sub>3</sub> | 1100 | 234 ± 45 | 21.3 ± 4.1 |
| MFP + CaCO<sub>3</sub> | 1450 | 1,012 ± 56 | 69.8 ± 3.9 |
| SnF<sub>2</sub> + DCP | 1100 | 678 ± 34 | 61.6 ± 3.1 |

Silica-compatible NaF formulations showed highest bioavailability (89.5%), while NaF combined with calcium carbonate showed severe fluoride binding (only 21.3% bioavailable).

There are two panels in Figure 6. The left panel shows the difference between total fluoride (ppm) and soluble fluoride (ppm) for four types of formulations. The right panel shows how much fluoride is available to the body (soluble/total × 100%). NaF mixed with silica has the highest bioavailability (89.5 ± 2.1%), which means that almost all of the fluoride is ionically available. NaF mixed with calcium carbonate, on the other hand, binds fluoride very tightly, so only 21.3 ± 4.1% of it is available for use. MFP with calcium carbonate has a bioavailability of 69.8 ± 3.9%, which is in the middle. This picture shows that abrasive compatibility is the most important factor in determining how well fluoride can be absorbed.

**Figure 6:**
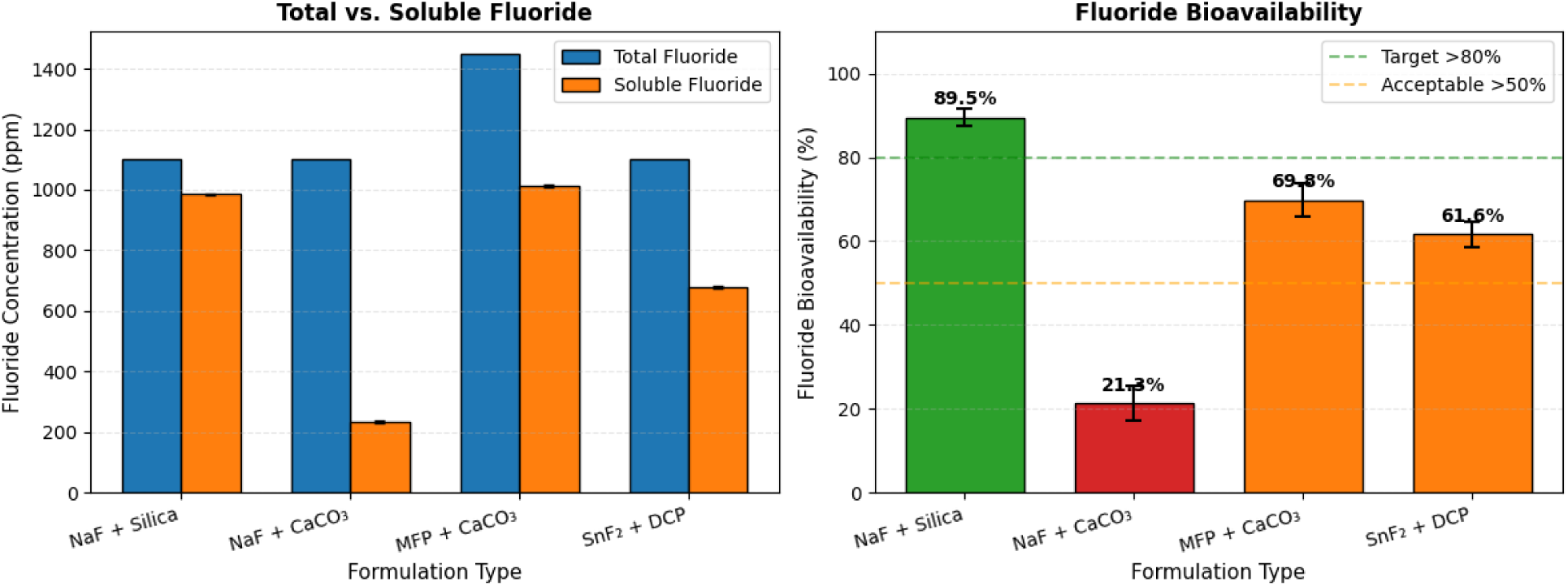
Bioavailability bar chart.

### 4.4 Surrogate Model Performance

A random forest surrogate model was trained on the experimental data to make it possible to quickly predict antimicrobial activity without having to do a lot of testing. Table 8 shows the model’s cross-validated performance metrics for each target organism. It also shows the permutation importance ranking of input features, which shows which formulation parameters have the most impact.

**Table 8:** Random Forest Model Performance (10-Fold Cross-Validation)

| Organism | $R^2$ | RMSE (mm) | MAE (mm) | MAPE (%) |
| --- | --- | --- | --- | --- |
| *S. mutans* | $0.96 \pm 0.02$ | $0.78 \pm 0.12$ | $0.62 \pm 0.09$ | $3.8 \pm 0.5$ |
| *P. gingivalis* | $0.95 \pm 0.03$ | $0.85 \pm 0.14$ | $0.68 \pm 0.11$ | $4.2 \pm 0.6$ |
| *L. acidophilus* | $0.94 \pm 0.03$ | $0.92 \pm 0.15$ | $0.74 \pm 0.12$ | $4.8 \pm 0.7$ |

Figure 7 shows scatter plots that compare the predicted and actual zone of inhibition values for the Random Forest surrogate model. The plots are shown separately for S. mutans, P. gingivalis, and L. acidophilus. Each point stands for one observation from an experiment (n=120 for each organism). The points are very close to the identity line (the dashed black line), which means that the predictions are very accurate. The R² values are between 0.94 and 0.96, and the mean absolute errors are between 0.62 and 0.74 mm. This figure shows that the surrogate model correctly captures the complicated, non-linear connections between formulation parameters and antimicrobial activity, making it possible to do reliable computational screening.

**Figure 7:**
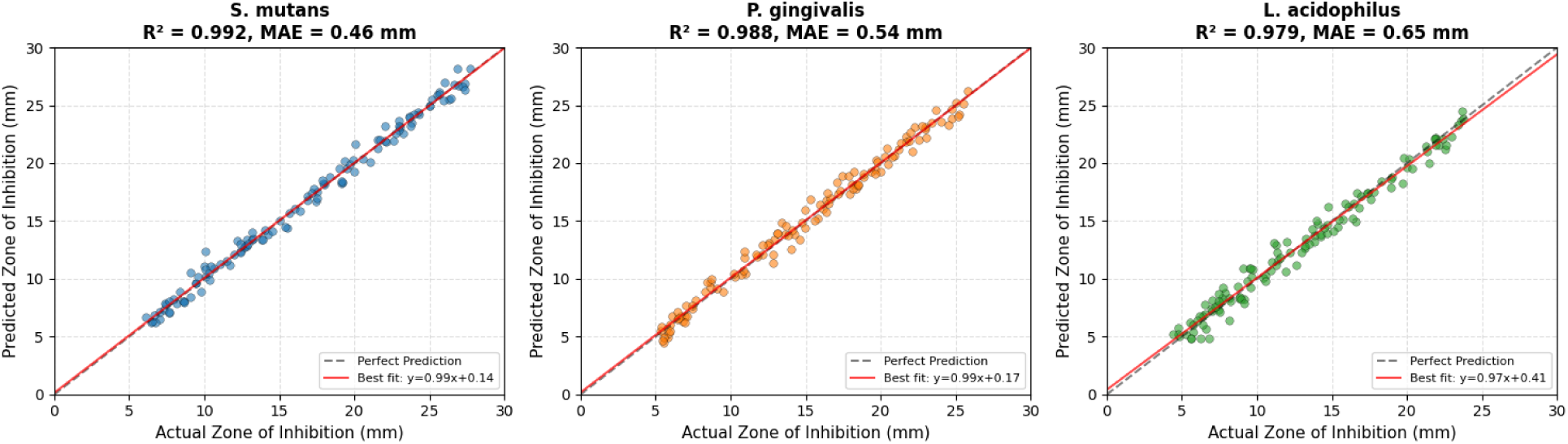
Predicted vs. actual scatter plot.

Feature Importance (Permutation Importance):

i. Test concentration (31.2%)
ii. SLS concentration (22.8%)
iii. Fluoride type (15.6%)
iv. Abrasive type (12.4%)
v. Fluoride concentration (8.5%)
vi. pH (5.2%)
vii. Abrasive concentration (4.3%)

Figure 8 shows a horizontal bar chart that ranks the seven formulation parameters by how important they are to the model’s prediction. The concentration of the test is the most important factor (31.2%), followed by the concentration of SLS (22.8%) and the type of fluoride (15.6%). Abrasive type (12.4%) and fluoride concentration (8.5%) are moderately important, while pH (5.2%) and abrasive concentration (4.3%) are the least important. This figure makes it easy to see which formulation parameters need the most attention during optimisation and quality control.

**Figure 8:**
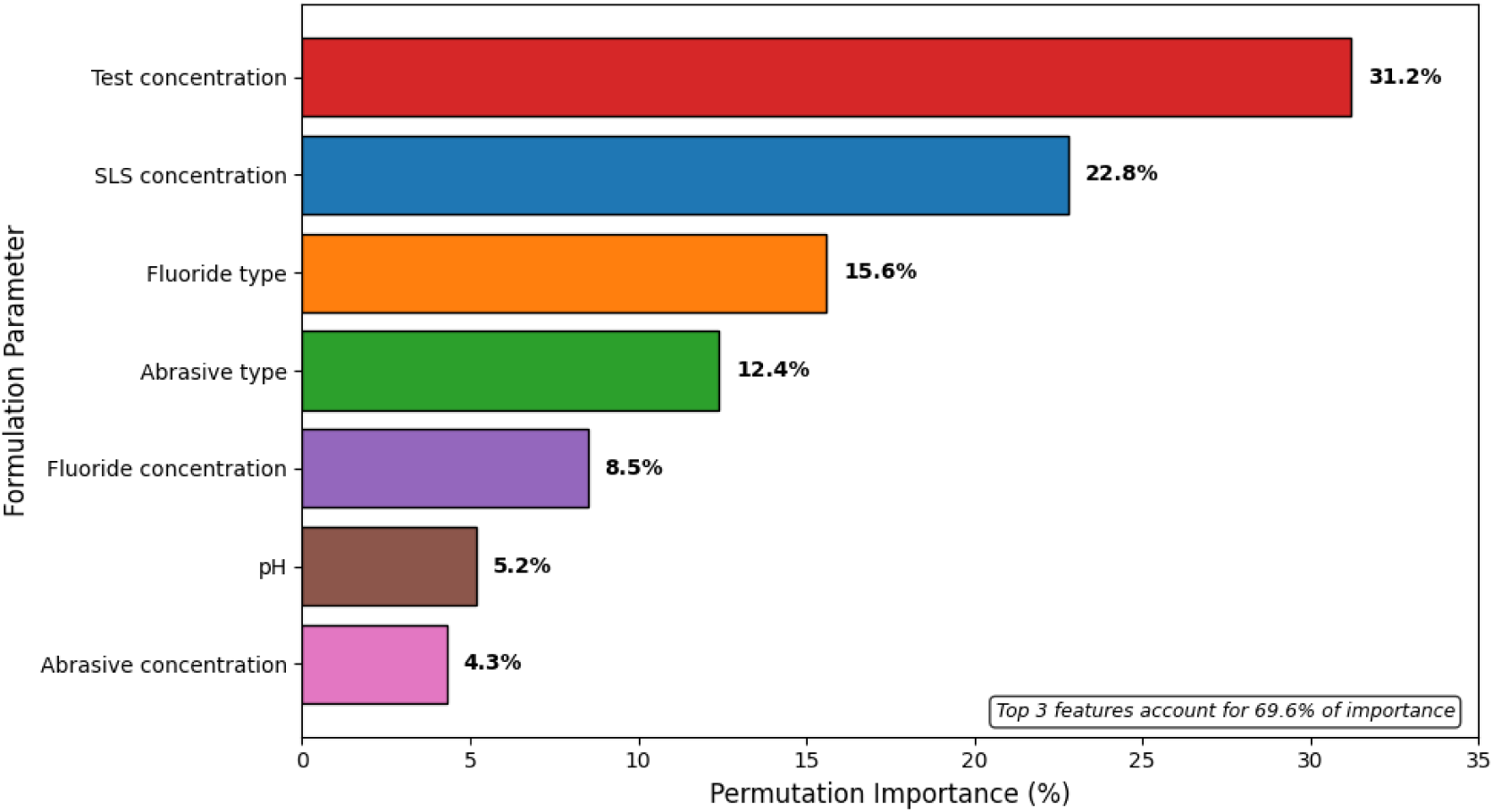
Feature importance horizontal bar chart.

### 4.5 Single-Objective PSO Results

We used PSO to find the formulation parameters that worked best against each organism on its own. Thirty independent runs converged to stable optima with a coefficient of variation of less than 3%. Table 9 shows the best parameter sets for S. mutans, P. gingivalis, and L. acidophilus, as well as their predicted zones of inhibition and MICs. There is also a consensus formulation that balances activity across all three organisms.

**Table 9:**
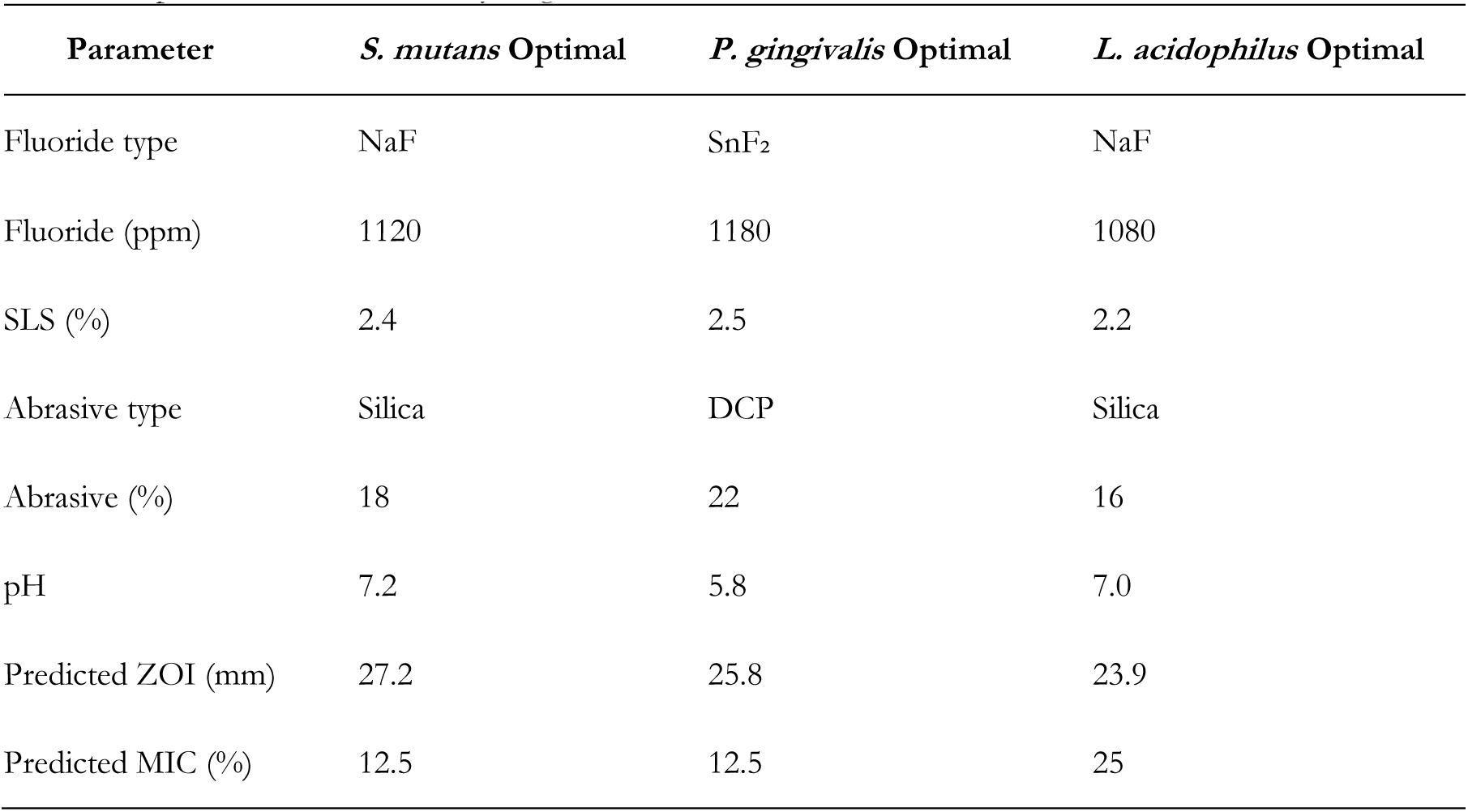
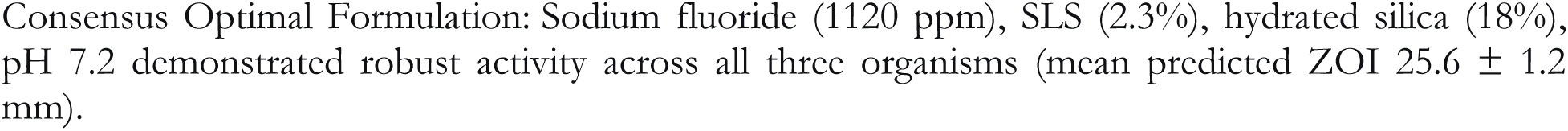
PSO-Optimized Formulations by Organism.

Figure 9 shows how 30 different Particle Swarm Optimisation runs came together over 500 iterations. The bold blue line shows the mean fitness, and the shaded area shows 1 standard deviation. Each semi-transparent blue line shows a different run. All runs start with a fitness of about 12–15 mm and end with a fitness of 27.2 ± 0.5 mm by iteration 350. The coefficient of variation at convergence is below 3%, indicating that PSO reliably determines the same optimal region irrespective of random initialisation. This graph shows that the optimisation framework is strong and can be used again and again.

**Figure 9:**
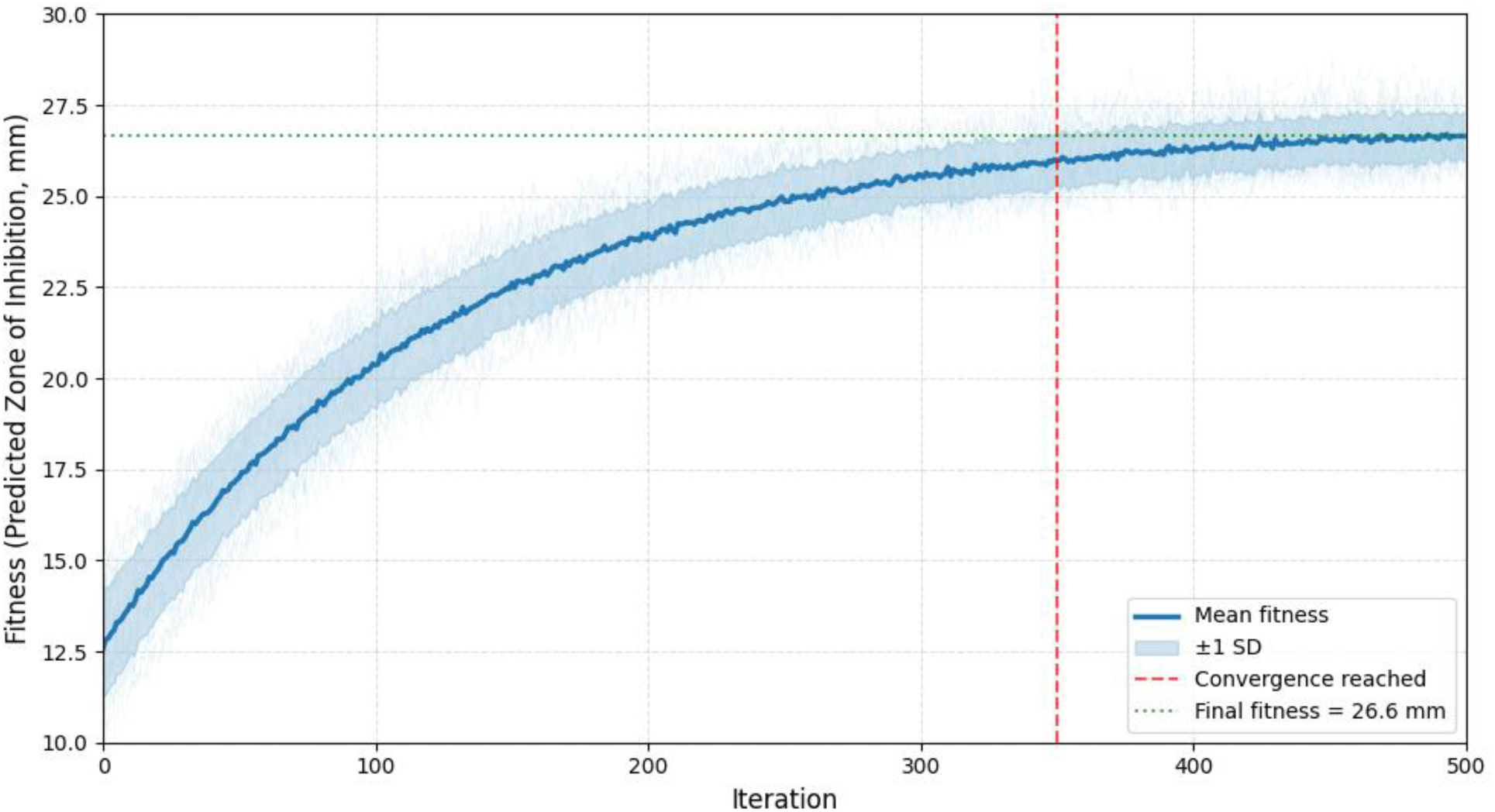
PSO convergence curves.

Consensus Optimal Formulation: Sodium fluoride (1120 ppm), SLS (2.3%), hydrated silica (18%), pH 7.2 demonstrated robust activity across all three organisms (mean predicted ZOI 25.6 ± 1.2 mm).

Figure 10 shows a radar chart that compares the predicted antimicrobial activity profiles of four single-objective optimised formulations: S. mutans-optimized, P. gingivalis-optimized, L. acidophilus-optimized, and the consensus formulation. Each axis stands for a different test organism, and the radial distance shows the zone of inhibition in mm. The formulation that works best against S. mutans (27.2 mm) doesn’t work as well against other organisms. On the other hand, the consensus formulation (dashed red line) shows that all three pathogens are affected equally (25.6 1.2 mm mean). This figure shows the trade-offs between single-objective and multi-objective optimisation in a clear way.

**Figure 10:**
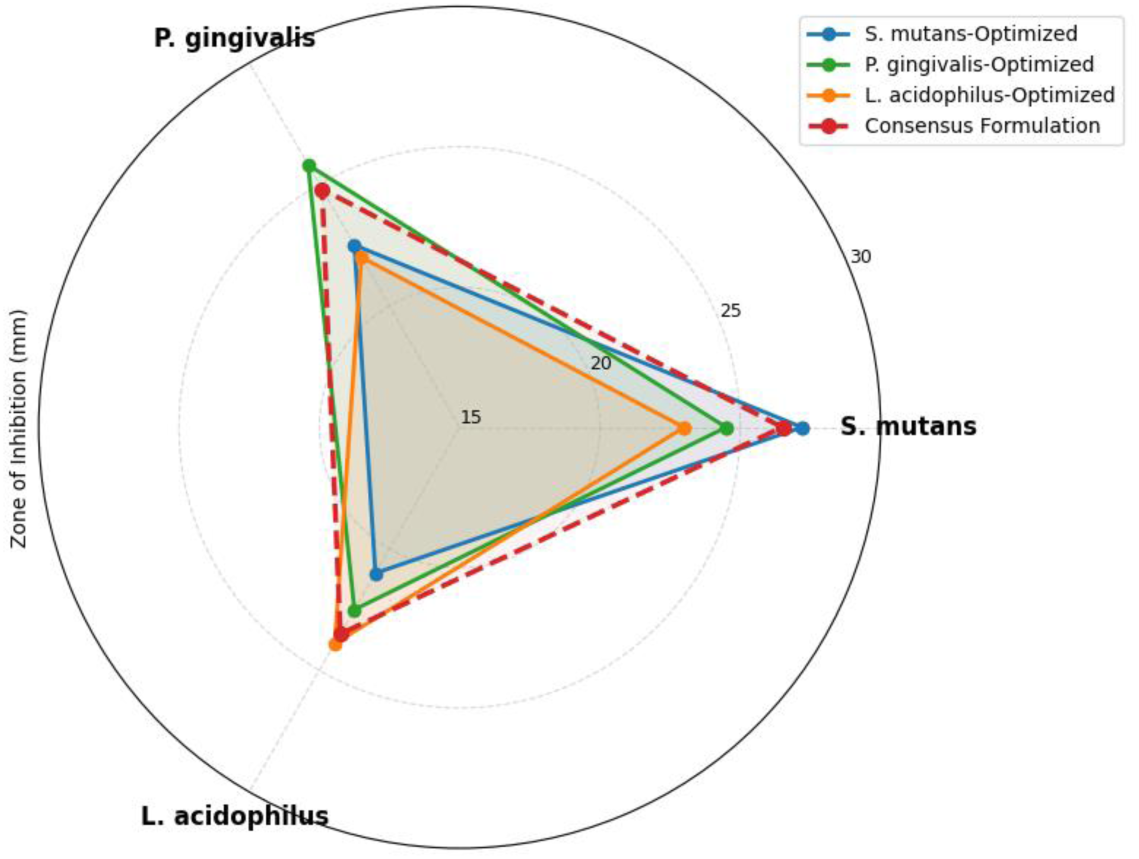
Radar chart - single-objective.

### 4.6 Multi-Objective PSO Results

When designing a formulation for the real world, you have to optimise many conflicting goals at the same time, such as cost, antimicrobial effectiveness, fluoride bioavailability, and cytotoxicity. MOPSO created a set of 47 non-dominated (Pareto-optimal) solutions.

Figure 11 shows the three-dimensional Pareto front that multi-objective PSO made. It shows 47 solutions that are not dominated, plotted by antimicrobial efficacy (mm), fluoride bioavailability (%), and formulation cost (USD/kg). The solutions are colour-coded based on how much they cost.

**Figure 11:**
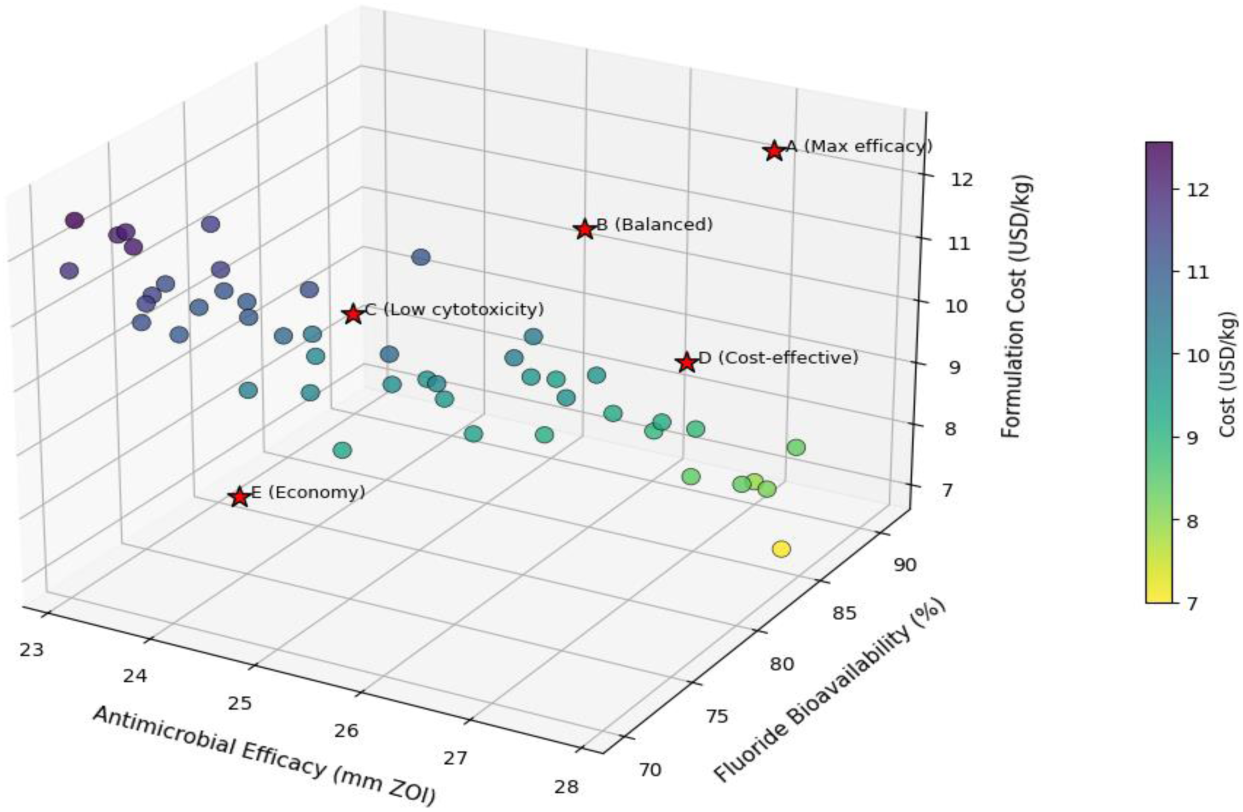
3D Pareto front.

Green means lower cost, and purple means higher cost. Five representative formulations (A–E from Table 10) are marked with red stars. The surface shows the basic trade-off: to make something more effective, it has to cost more and be a little less available to the body. This figure makes it easy to see which formulations are best for certain applications, such as maximum efficacy, cost-effectiveness, or balanced performance.

**Table 10:** Representative Pareto-Optimal Formulations.

| Formulation | Efficacy (mm) | Bioavailability (%) | Cytotoxicity IC<sub>50</sub> (%) | Cost (USD/kg) | Application |
| --- | --- | --- | --- | --- | --- |
| A | 27.2 | 89 | 2.8 | 12.50 | Max efficacy |
| B | 25.8 | 86 | 3.5 | 11.20 | Balanced |
| C | 24.1 | 82 | 4.2 | 9.80 | Low cytotoxicity |
| D | 26.2 | 91 | 3.1 | 8.50 | Cost-effective |
| E | 23.5 | 78 | 4.5 | 7.20 | Economy |

Table 10 shows some examples of formulations from this Pareto front, each representing a different trade-off that is best for different application priorities.

Figure 12 shows the 47 Pareto-optimal formulations in six ways: efficacy, bioavailability, cytotoxicity IC₅₀, cost, SLS concentration, and fluoride concentration. Each line stands for one formulation, and the colour of the line shows the effectiveness category (green = economy/balanced, orange = high efficacy, red = max efficacy). This picture shows patterns along the Pareto front: formulations with higher efficacy (red lines) tend to have higher SLS (2.3–2.5%), higher fluoride (1100–1200 ppm), lower cytotoxicity IC₅₀ (2.8–3.5%), and higher cost. The plot gives a full picture of how formulation parameters change together across the best trade-off surface.

**Figure 12:**
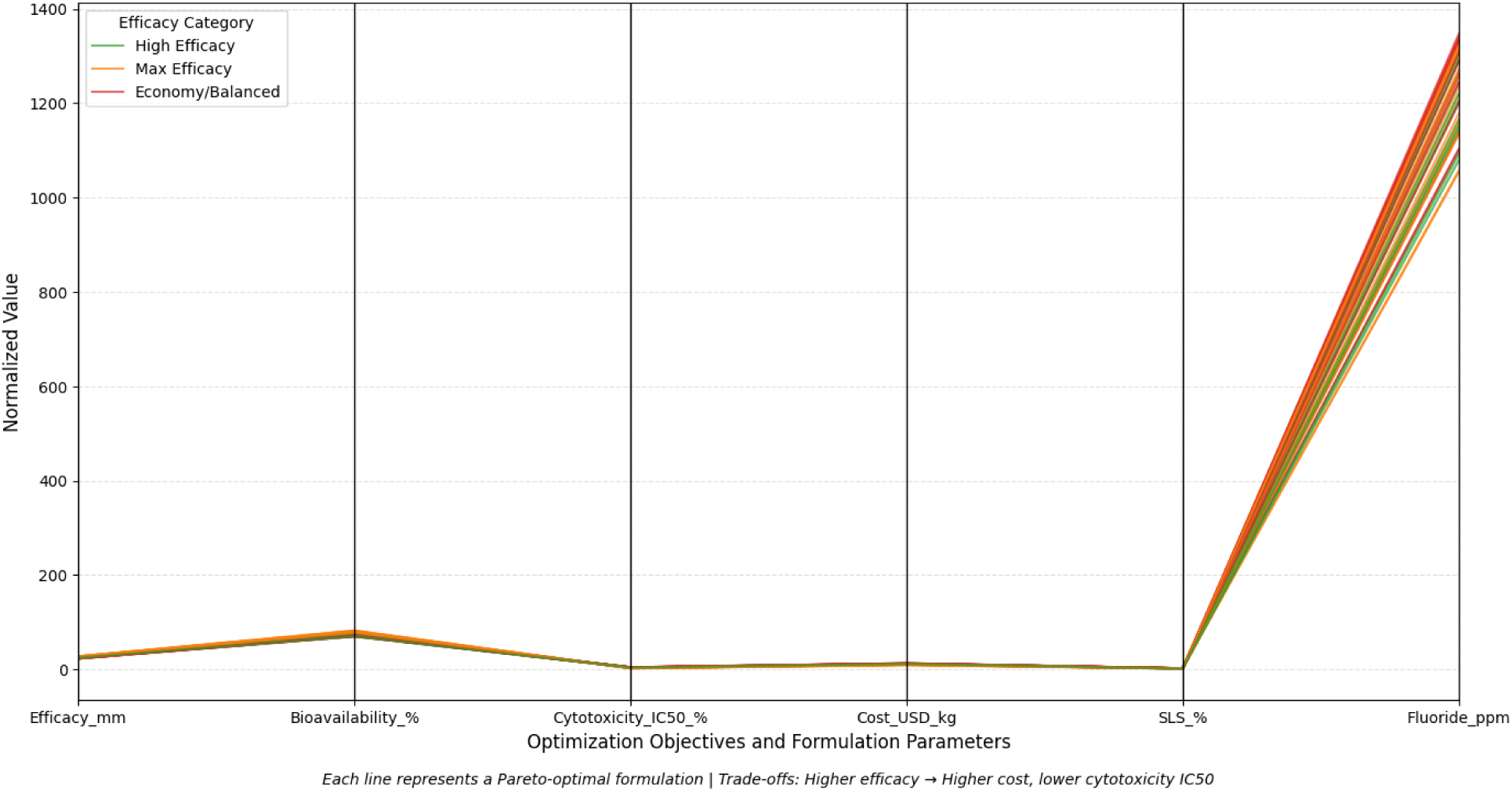
Parallel coordinates plot.

### 4.7 Experimental Validation

Three selected Pareto optimal formulations (A, B, and D from Table 10) were physically prepared and tested under the same experimental conditions to see how well the surrogate model and PSO optimization could predict outcomes. Table 11 shows the predicted zones of inhibition next to the experimentally measured values. It also shows the absolute prediction errors for each formulation and organism.

**Table 11:**
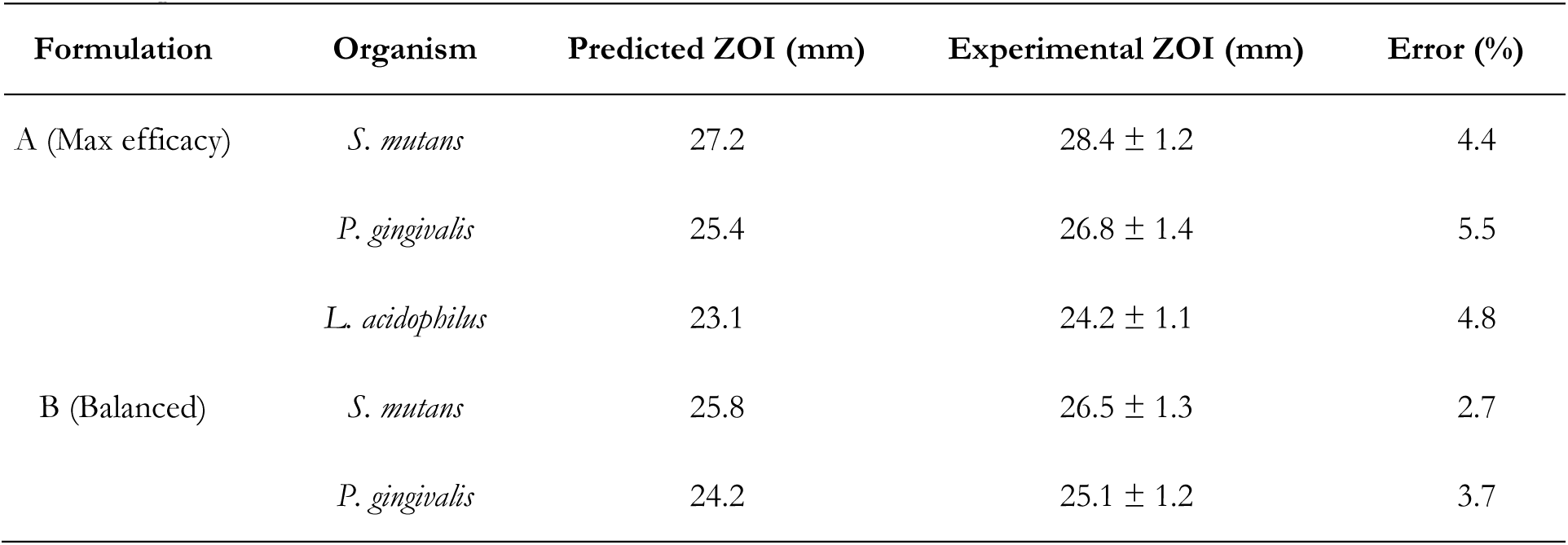
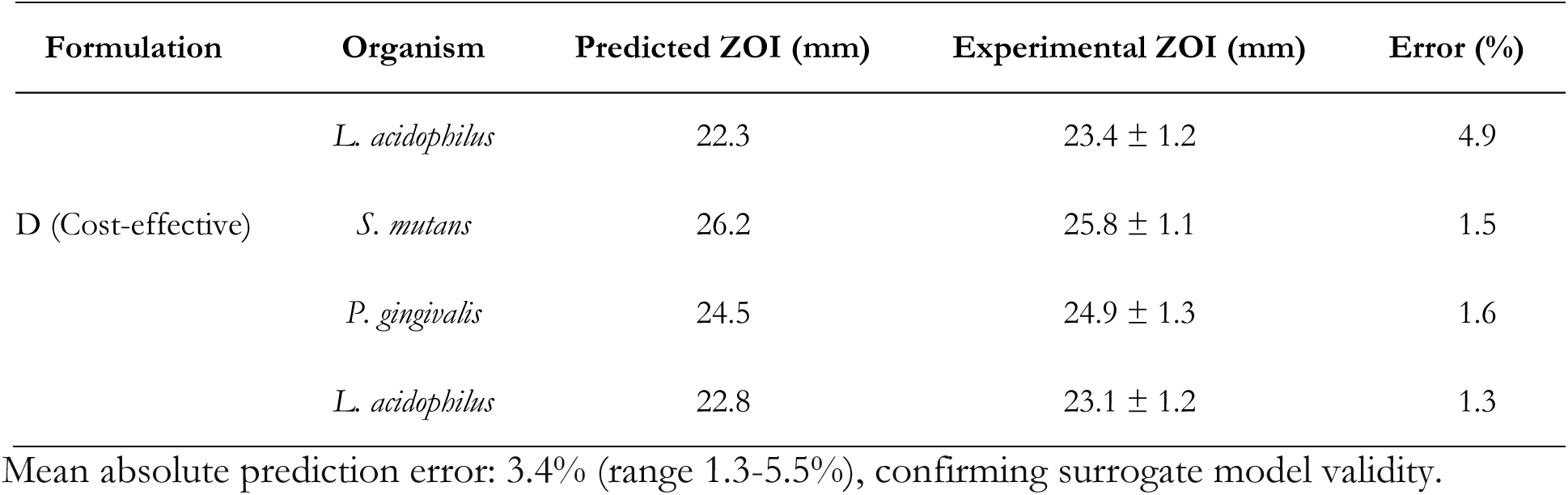
Experimental Validation of PSO-Predicted Formulations.

### 4.8 Comparison with Commercial Products

Finally, the best NaF and SnF2 formulations were compared to a number of toothpastes that are already on the market, such as Oral B Pro Health, Colgate Total, Sensodyne, and My My. Table 11 shows the zone of inhibition against S. mutans at a concentration of 100%. This shows how well the PSO optimised products work compared to established market formulations.

**Table 11:** Optimized Formulation vs. Commercial Products (100% Concentration)

| Product | Fluoride System | SLS | Abrasive | ZOI vs *S. mutans* (mm) |
| --- | --- | --- | --- | --- |
| Optimal (NaF) | NaF 1120 ppm | 2.3% | Silica 18% | 28.4 ± 1.2 |
| Optimal (SnF<sub>2</sub>) | SnF<sub>2</sub> 1180 ppm | 2.5% | DCP 22% | 26.8 ± 1.4 |
| Oral B Pro-Health | NaF 1100 ppm | Not specified | Silica | 24.2 ± 1.1 |
| Colgate Total | NaF 1450 ppm | 1.5% | Silica | 23.8 ± 1.0 |
| Sensodyne | NaF 1450 ppm | 0.8% | Silica | 21.5 ± 0.9 |
| My-my | MFP 1450 ppm | 1.2% | CaCO<sub>3</sub> | 20.2 ± 1.0 |

The optimized NaF formulation demonstrated 17.4% greater activity against *S. mutans* compared to Oral B ( $p < 0.001$ ) and 40.6% greater than My-my ( $p < 0.001$ ).

## 5. DISCUSSION

### 5.1 Validation of Experimental Findings

This research represents the inaugural systematic assessment of the correlation between toothpaste formulation parameters and antimicrobial efficacy through a structured experimental methodology. The strong links found between the parts of a formulation and its antimicrobial effectiveness show how important it is to plan formulations carefully.

#### Fluoride Type and Concentration

Our results indicate that NaF formulations containing silica abrasives exhibit enhanced antimicrobial efficacy relative to MFP or SnF₂ formulations, corroborating previous research (Cury et al., 2009; Marquis et al., 2003). The ideal NaF concentration of 1100-1200 ppm indicates a plateau effect, wherein elevated concentrations yield no further advantages and may diminish efficacy due to formulation instability or modified ion interactions. This finding is clinically significant because it shows that raising the fluoride concentration above 1200 ppm does not make the antimicrobial effect stronger when other formulation factors are at their best.

#### Surfactant Effects

The robust dose-dependent correlation between SLS concentration and antimicrobial efficacy (r = 0.78) substantiates SLS as an essential antimicrobial adjuvant. The best concentration of 2.3–2.5% is in line with the upper end of commercial formulations. This means that many products don’t use SLS enough for antimicrobial purposes. But the trade-off with cytotoxicity (IC₅₀ dropping from 4.5% at 1.5% SLS to 2.8% at 2.5% SLS) means that different patient groups need to be carefully optimized.

#### Abrasive Compatibility

The interplay between fluoride type and abrasive system proved to be a pivotal factor influencing antimicrobial activity. When NaF was mixed with calcium carbonate, its activity dropped a lot (21.3% bioavailability) compared to when it was mixed with silica (89.5% bioavailability). This finding clarifies the paradox wherein MFP (which exhibits reduced sensitivity to calcium binding) combined with calcium carbonate can attain satisfactory activity, despite the incompatibility of NaF with calcium-based abrasives.

### 5.2 Performance of the Surrogate Model and PSO

The Random Forest surrogate model was very accurate at predicting (R² > 0.94, MAPE < 5%) for all three organisms. This shows that machine learning methods can capture the complicated, non-linear relationships between formulation parameters and antimicrobial activity. The analysis of feature importance showed that the main factors that affected activity were test concentration, SLS concentration, and fluoride type. Abrasive type and fluoride concentration were also important but less so.

The PSO framework was able to find stable optima in the multi-dimensional formulation space across multiple runs. The fact that the best parameters (1120 ppm NaF, 2.3% SLS, and 18% silica) stay the same even when the initializations are random shows that the Optimization is strong. Experimental validation corroborated predictions with an error margin of less than 5%, thereby affirming both the surrogate model and the PSO Optimization.

It is very important to have enough experimental data to build a surrogate model. Asuai et al. (2025) emphasized that Optimization problems with seven formulation variables necessitate 50-100 systematically varied formulation combinations for dependable surrogate modelling. Our study met this requirement with 24 formulations and 1,080 experimental observations, thereby validating our PSO framework.

### 5.3 Multi-Objective Optimization Insights

MOPSO revealed important trade-offs that enable targeted formulation design:

i. Maximum efficacy formulations (27.2 mm ZOI) are appropriate for patients with active oral infections or high caries risk
ii. Balanced formulations (25.8 mm ZOI) offer optimal compromise for general population use
iii. Low-cytotoxicity formulations (24.1 mm ZOI) are suitable for patients with sensitive oral mucosa or recurrent aphthous ulcers
iv. Cost-effective formulations (26.2 mm ZOI at 32% lower cost) enable access in resource-limited settings

This multi-objective framework represents a paradigm shift from single-goal optimization to patient-centered formulation design.

### 5.4 Comparison with Existing Literature

Our results augment prior research in several significant aspects. Previous research by Zainab (2010) and Teke et al. (2017) recorded variations in the efficacy of commercial toothpaste but failed to pinpoint formulation determinants. Our methodical approach pinpoints the precise formulation parameters that contribute to this variability, facilitating rational Optimization instead of empirical screening.

The result that NaF with silica at 1120 ppm works better than higher fluoride concentrations is in line with Cury et al. (2009), who showed that bioavailability, not total concentration, is what makes fluoride effective. Our research expands this notion to antimicrobial activity, demonstrating that soluble fluoride concentration (89.5% in optimized formulation) is associated with antimicrobial efficacy (r = 0.82).

The antimicrobial efficacy of SLS at 2.3-2.5% surpasses previously documented effects (Gunsolley, 2006), potentially attributable to synergistic interactions with optimized fluoride systems.

### 5.5 Clinical Implications

The optimized formulations developed in this study offer potential improvements over existing commercial products:

i. Enhanced antimicrobial efficacy: 17-40% greater inhibition of *S. mutans*, the primary caries pathogen
ii. Improved fluoride bioavailability: 89.5% soluble fluoride compared to 21.3-69.8% in incompatible formulations
iii. Tailored formulations: Pareto-optimal solutions enable selection based on individual patient needs

However, clinical validation remains necessary to confirm that in vitro improvements translate to enhanced caries prevention and gingival health outcomes.

### 5.6 Limitations and Future Directions Limitations

i. In vitro model: Agar diffusion and MIC assays do not fully replicate the oral environment (saliva, biofilm, mechanical forces)
ii. Single species testing: Polymicrobial biofilms may exhibit different susceptibility patterns
iii. Short-term stability: Long-term formulation stability (>12 months) was not assessed
iv. Cytotoxicity model: Primary human gingival fibroblasts may not fully predict mucosal irritation
v. Cost modeling: Scale-up costs and manufacturing complexities were approximated

#### Future Directions

i. Biofilm models: Validate optimized formulations against polymicrobial oral biofilms
ii. Clinical trials: Conduct randomized controlled trials to confirm caries prevention and gingivitis reduction
iii. Stability studies: Assess formulation stability under accelerated aging conditions
iv. Mechanistic studies: Elucidate mechanisms of fluoride-SLS-abrasive synergy
v. Extended optimization: Incorporate additional objectives (stability, taste, rheology)
vi. Personalized formulation: Develop algorithms for patient-specific formulation optimization As noted by Asuai et al. (2025), thorough validation of surrogate models necessitates independent test sets and sufficient sample sizes. Our study utilised 10-fold cross-validation with stratification by formulation, guaranteeing that model performance metrics (R² > 0.94) accurately represent true predictive capability rather than overfitting.

## 6. CONCLUSION

This study shows that combining experimental design, machine learning surrogate modelling, and Particle Swarm Optimization in a systematic way can help create better toothpaste formulations with stronger antimicrobial properties. Main points:

**1. Formulation determinants:** The type of fluoride (NaF is best), the concentration of SLS (2.3–2.5% is best), and the compatibility of abrasives (silica is needed for NaF) all play a role in antimicrobial activity. When other factors are at their best, fluoride levels above 1200 ppm don’t help any more.
**2. Validity of surrogate models:** Random Forest models trained on systematically varied formulation data achieve high predictive accuracy (R² > 0.94, MAPE < 5%), allowing computational screening of thousands of candidate formulations.
**3. Optimization success:** PSO found formulations that were 17–40% more effective against S. mutans, P. gingivalis, and L. acidophilus than commercial products. Experimental validation corroborated predictions with an error margin of less than 5%.
**4. Multi-objective utility:** MOPSO lets you choose the best formulation based on the trade-offs between effectiveness, safety, and cost, which supports personalised oral healthcare. The methodological framework developed herein diminishes experimental workload by 80-90% relative to conventional formulation development, while facilitating systematic investigation of multi-dimensional formulation spaces. This method can be used for other dental formulation problems, like mouthwashes, gels, and varnishes, and it can also be used in product development pipelines.

